# Modeling heterogeneity in single-cell perturbation states enhances detection of response eQTLs

**DOI:** 10.1101/2024.02.20.581100

**Authors:** Cristian Valencia, Aparna Nathan, Joyce B. Kang, Laurie Rumker, Hyunsun Lee, Soumya Raychaudhuri

**Affiliations:** Center for Data Sciences, Brigham and Women’s Hospital, Boston, MA, USA; Division of Genetics, Department of Medicine, Brigham and Women’s Hospital and Harvard Medical School, Boston, MA, USA; Division of Rheumatology, Inflammation, and Immunity, Department of Medicine, Brigham and Women’s Hospital and Harvard Medical School, Boston, MA, USA; Department of Biomedical Informatics, Harvard Medical School, Boston, MA, USA; Program in Medical and Population Genetics, Broad Institute of MIT and Harvard, Cambridge, MA, USA; Immune Tolerance Network, Boston, MA, USA

## Abstract

Identifying response expression quantitative trait loci (reQTLs) can help to elucidate mechanisms of disease associations. Typically, such studies model the effect of perturbation as discrete conditions. However, perturbation experiments usually affect perturbed cells heterogeneously. We demonstrated that modeling of per-cell perturbation state enhances power to detect reQTLs. We use public single-cell peripheral blood mononuclear cell (PBMC) data, to study the effect of perturbations with *Influenza A virus* (IAV), *Candida albicans* (CA), *Pseudomonas aeruginosa* (PA), and *Mycobacterium tuberculosis* (MTB) on gene regulation. We found on average 36.9% more reQTLs by accounting for single cell heterogeneity compared to the standard discrete reQTL model. For example, we detected a decrease in the eQTL effect of rs11721168 for *PXK* in IAV. Furthermore, we found that on average of 25% reQTLs have cell-type-specific effects. For example, in IAV the increase of the eQTL effect of rs10774671 for *OAS1* was stronger in CD4+T and B cells. Similarly, in all four perturbation experiments, the reQTL effect for *RPS26* was stronger in B cells. Our work provides a general model for more accurate reQTL identification and underscores the value of modeling cell-level variation.

## Introduction

Genome wide association studies (GWAS) have elucidated loci underpinning human complex traits, including autoimmune diseases such as rheumatoid arthritis (RA)^1^ and systemic lupus erythematosus (SLE)^2^, and infection-related traits such as HIV control^3^ and COVID-19 susceptibility^4^. However, the mechanisms through which the risk alleles within these loci act remains elusive. One possibility is that alleles alter the regulation of gene expression. Mapping expression quantitative trait loci (eQTLs) aims to improve the interpretation of GWAS results by linking genetic variants to changes in the expression levels of specific genes. Efforts such as the Genotype-Tissue Expression (GTEx) Project and the Database of Immune Cell Expression, Expression quantitative trait loci and Epigenomics (DICE)^5^ have identified thousands of eQTLs from healthy individuals using aggregated expression profiles from multiple cell types (bulk transcriptomics). However, eQTLs can be context-dependent and have different effects in specific contexts, for example in different cell states or in different physiologic conditions. Previous studies noted that less than 10% of the eQTL effects found at homeostasis colocalize with GWAS loci^6^. One possibility is that the regulatory effect of disease alleles may be seen only in very specific cellular contexts that are critical to disease initiation^7^.

One way to understand context-dependent gene regulation is by studying response eQTLs (reQTLs): genetic variants associated with the change in gene expression after environmental perturbations. reQTL studies typically divide samples from each individual into multiple conditions—e.g. external stimulations—and map the reQTL by measuring the effects of both genotype and genotype (G) by environment (E) interactions (G_x_E) on bulk gene expression profiles. This approach is critical to understand how genetic variation may act in disease-specific conditions that are missed in typical eQTL studies, and it has previously helped link disease-associated loci to condition-specific regulatory variation. For example, to elucidate the genetic mechanisms of autoimmune diseases, previous efforts studied reQTLs in putative causal immune cell types such as monocytes and dendritic cells perturbed by different disease-relevant stimuli; these stimuli included cytokines, pathogenic antigens, or T cell receptor stimulation^8–11^. However, in practice ex vivo experimental conditions affect cells heterogeneously^12–14^. For example, during T cell stimulation, factors such cell density or endogenous variability on protein levels can resulted in different activation states^15,16^.

Current models to find reQTLs cannot disentangle heterogeneous response eQTLs effects across different perturbation states. Accounting for this heterogeneity in statistical models may improve powered to detect re-eQTLs.

Single cell RNA-seq offers the opportunity to infer the response state of each cell from its transcriptional profile, revealing the heterogeneity of cellular response upon perturbation. We and others have recently demonstrated that mapping eQTLs at single cell resolution can identify heterogeneous eQTL effects along continuous cell-states^17,18^. Recently, a non-linear model was used to estimate G_x_E effects at cell resolution along continuous trajectories of innate antiviral response in iPSC-derived fibroblasts^19^.

However, reQTL models typically only offer a single format to represent perturbation response^20,21^, which neglects the diversity of potential forms of perturbation response. We need frameworks that effectively incorporated the heterogeneity of the response to perturbation in the analysis of reQTLs at single cell resolution. We hypothesize that effective modeling of per-cell perturbation state using linear models may enhance power to detect response eQTLs.

Here, we demonstrated that modeling eQTLs effects in single cells and scoring cells along a continuous perturbation score improves power to detect response eQTLs relative to reQTL models with a binary representation of the perturbation state.

Furthermore, we demonstrated that our approach could help to prioritize genetic effects in disease-relevant contexts.

## Results

### General framework

Our framework to map reQTLs accounts for heterogeneity in cell perturbation; we hypothesize that this will increase power to detect reQTLs (**Figure 1a**). First, we define a continuous score to quantify the degree of perturbation response of each cell. The underlying assumption is that perturbation may induce changes in the transcriptional profile of a cell that are proportional to the degree of response of the cell. To systematically identify axes of variation in the transcriptional profile of perturbed and non-perturbed cells, we use principal components analysis (PCA) and correct technical or biological effects with Harmony to obtain corrected expression PCs (hPCs) (**METHODS**). We assume that one or multiple hPCs may capture the effect of perturbation in a cell. To define a continuous perturbation score, we used a penalized logistic regression with hPCs as independent variables to predict the log odds of being in the pool of perturbed cells, which serves as a surrogated of the cell’s degree of response to the experimental perturbation. We use this approach instead of focusing on pre-defined gene sets like the interferon-stimulated gene pathway for viral perturbations, which may not accurately represent the changes on the transcriptome in specific tissues, cells or perturbation experiments. Then, to find response eQTLs, we used a Poisson mixed effects model (PME) of gene expression in single cells as a function of genotype and genotype interactions with discrete perturbation and the independent effect of the continuous perturbation states, accounting for confounders and batch as we previously described^17^ (**METHODS**). We calculated the independent effect of the continuous perturbation state by first regressing out the discrete perturbation effect and then using the residuals in the PME model (METHODS).

**Figure 1.**
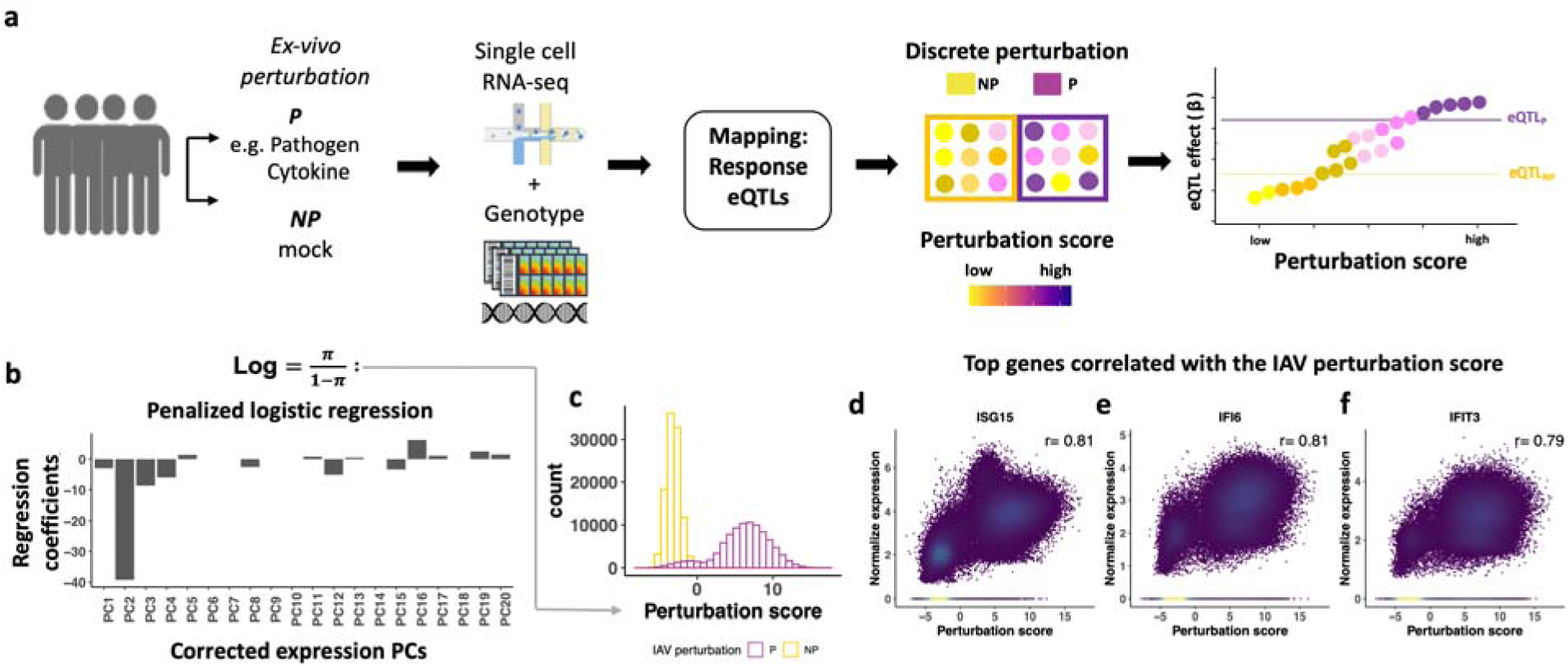
Framework to identify response eQTL at single-cell resolution. **(a)** Cells from each donor are collected and divided into the non-perturbed (NP) and perturbed (P) conditions. Then, the transcriptional profile of each cell is ascertained using single cell RNA-seq. To identify expression quantitative trait loci associated with the change in gene expression after perturbations (reQTL), we used a Poisson mixed effects model of gene expression in single cells (**METHODS**). **(b)** The continuous perturbation score is based on penalized logistic regression that predicts discrete perturbation state based on the corrected expression PCs. This perturbation score can be used as a surrogate of the continuous response state of each cell. The plot shows the regression coefficients of the penalized logistic regression in the IAV perturbation. **(c)** Histogram showing the distribution of the IAV perturbation score color by the experimental condition. **(d-f)** Scatter plots showing correlation of the IAV perturbation score and the expression of top three correlated genes colored by cell density. *ISG15*=ISG15 ubiquitin like modifier, *IFI6*= interferon alpha inducible protein 6, *IFIT3*=interferon induced protein with tetratricopeptide repeats 3.

### Quantifying perturbation states

We applied our framework to study the effect of ex vivo stimulation with *Influenza A virus* (IAV), *Candida albicans* (CA), *Pseudomonas aeruginosa* (PA), and *Mycobacterium tuberculosis* (MTB) on gene regulation. We used transcriptional profiles from 299,879 peripheral blood mononuclear cells (PBMC) from 89 donors previously published in Randolph et al. (Science, 2022)^22^ and 475,333 PBMCs from 120 donors in Oelen et al (Nature Comm, 2022^21^) (**METHODS**). We expected that each perturbation may have different effect on cell states, so we defined perturbation scores independently for each stimulation condition (**Figure 1 b-c and Figure S1 a-c**) using a penalization factor with minimal cross-validation error (**Figure S1 d-g**).

We found that the perturbation score corresponds to changes in single cell transcriptional profiles. For example, the IAV perturbation score correlates with the expression of genes such as *ISG15* (Pearson r=0.81, **Figure 1 d**), *IFI6* (Pearson r=0.81, **Figure 1 e**), and *IFIT3* (Pearson r=0.79, **Figure 1 f**) and these top correlated genes were enriched in the "interferon alpha response" MSigDB gene set^23,24^, (GSEA P adjusted=0.003) (**Table 1**, **Figure S2**). (gene set enrichment analysis GSEA^25^, **METHODS**). This is consistent with the characterized role of the type I interferon response during RNA-virus infection^26^. Similarly, the CA perturbation score correlates with the expression of genes such as *STAT1*, (Pearson r=0.50, **Figure 2 c**), *GBP1* (Pearson r=0.45, **Figure 2 d**), and *GBP5* (Pearson r=0.41, **Figure 2 e**), and these genes were enriched in the “interferon gamma response” MSigDB gene set (GSEA P adjusted=0.003) (**Table 1**, **Figure S2**). The PA perturbation score showed enrichment for genes in the “glycolysis” MSigDB gene set (GSEA P adjusted=0.02) (**Table 1**, **Figure S2**) such as *SDC2* (Pearson r=0.39, **Figure 2 c**) and genes from the “complement pathway” MSigDB gene set (GSEA P adjusted=0.09) (**Table 1**, **Figure S2**. like *MMP14* (Pearson r=0.43, **Figure 2 c**), and *CXCL1* (Pearson r=0.37, **Figure 2 c**).

**Figure 2.**
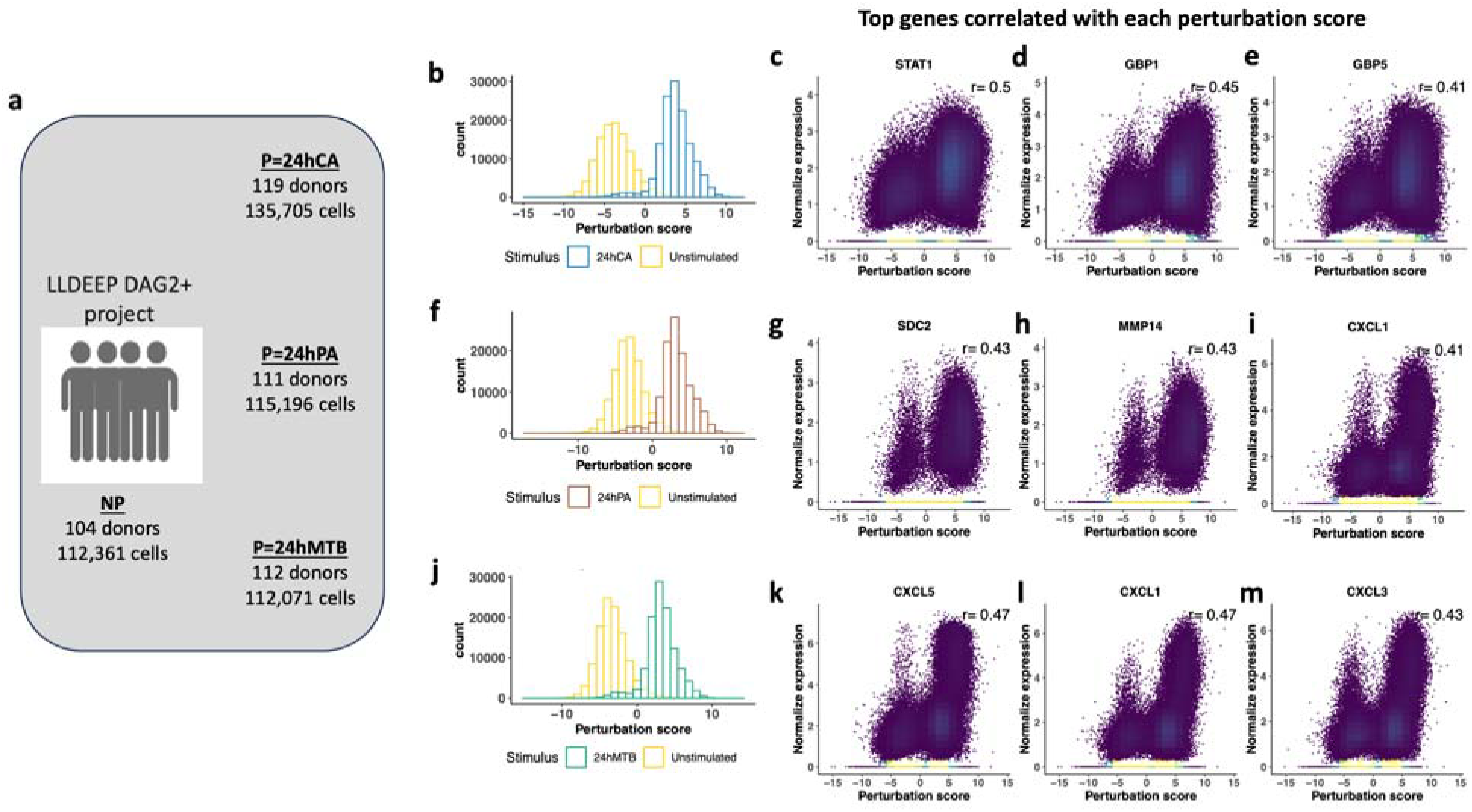
(a) Schematic of the perturbation score in the perturbation experiments from The Lifelines DEEP DAG2+ Project. **(a)** We obtained genotype and single cell mRNA data from 120 participants of The LifeLines DEEP prospective cohort (LLDEEP DAG2+ project). A total of 112,362 cells from 104 donors that were subject to non-perturbation conditions were included in the presented analysis. Similarly, we include cells perturbed with *Candida albicans* (CA), *Pseudomonas aeruginosa* (PA), and *Mycobacterium tuberculosis* (MTB) for 24 hours. Histogram showing the distribution of the perturbation score in the **(b)** CA, (f) PA and (j) MTB perturbation experiment. Scatter plots showing the correlation of the perturbation score and the expression of top correlated genes colored by cell density in the CA **(c-e)**, PA **(g-i)** and MTB **(k-m)** perturbation experiments. *STAT1*= signal transducer and activator of transcription 1, *GBP1*= guanylate binding protein 1, *GBP5*= guanylate binding protein 1, *RAB13*= RAB13, member RAS oncogene family, *CXCL5*= C-X-C motif chemokine ligand 5, *MMP14*= matrix metallopeptidase 14, *CXCL1*= C-X-C motif chemokine ligand 1, *CXCL3*= C-X-C motif chemokine ligand 3.

Furthermore, the genes such as *CXCL5* (Pearson=0.47, **Figure 2 c**), *CXCL1* (Pearson=0.47, **Figure 2 c**), *CXCL3* (Pearson=0.43, **Figure 2 c**) were the top 3 with expression that correlated with the MTB perturbation score and these genes were enriched for the “inflammatory response pathway” MSigDB gene set (GSEA P adjusted=0.02) (**Table 1**, **Figure S2**).

### Mapping response eQTLs

We decided to prioritize a set of eQTL with robust main effects. We first mapped eQTLs in a dataset of expression profiles including all cell types and perturbation conditions (NP=non-perturbed, P=perturbed). To find these main effect eQTLs in each perturbation experiment, we first generated pseudobulk profiles and identified the lead eQTL for each gene (**METHODS**). Furthermore, recognizing that a Poisson model may not be appropriate for some genes with high differential expression across cell states, we removed genes with inflated p values under permutation testing (**METHODS**); on average we removed 7.23 % of genes. We prioritized 1,892 eQTLs for IAV, 1,863 eQTLs for CA, 1,794 eQTLs for PA and 1,355 eQTLs for MTB data-sets. In a secondary analysis of the CA, PA and MTB experiments, we used 1,825 eQTLs that were significant (FDR-corrected p<0.05) in the original analysis by Oelen et al^21^ (**METHODS**).

Next, we tested which of these eQTLs with significant main effects varied in effect size after perturbation. To do this we modeled the gene expression in single cells as a function of genotype (G) and the interaction with the discrete perturbation state (G_x_Discrete) and the independent effect of the continuous perturbation state (G_x_Score) accounting for confounders and batch (**METHODS**) using the PME model described by Nathan et al 2022^17^. We assessed the significance of both G_x_Discrete and G_x_Score terms compared to a null model with no interactions using a two degrees-of-freedom likelihood ratio test (LRT), hereafter referred to as the 2df-model. We defined the total eQTL effect in each cell as the sum of the main eQTL effect (G) and the interaction effects (the sum of each interaction term multiplied by its corresponding perturbation state (NP=0, P=1; continuous perturbation state=perturbation score) (**METHODS**).

By using the 2df-model we found 166 reQTL in IAV, 770 reQTL in CA, 646 reQTL in MTB, and 594 reQTL in PA (LRT Q value <0.05, **Figure 3 a-d. Table S2-S5**). To confirm that the reQTLs are not spurious associations, we performed 500 permutations for each reQTL by randomly shuffling the perturbation state of the cell within each donor and excluded any reQTL with permutation p>0.05 (**METHODS**) (IAV=0 **Table S2**, CA=129 **Table S3**, PA=59 **Table S4**, and MTB=184 **Table S5**). We also compared the 2df-model results with a negative binomial model, which models overdispersion and observed an average correlation of 0.90 between the LRT P values of the two models (**Figure S4 a-d**), indicating that the p values from the PME model were not inflated.

**Figure 3.**
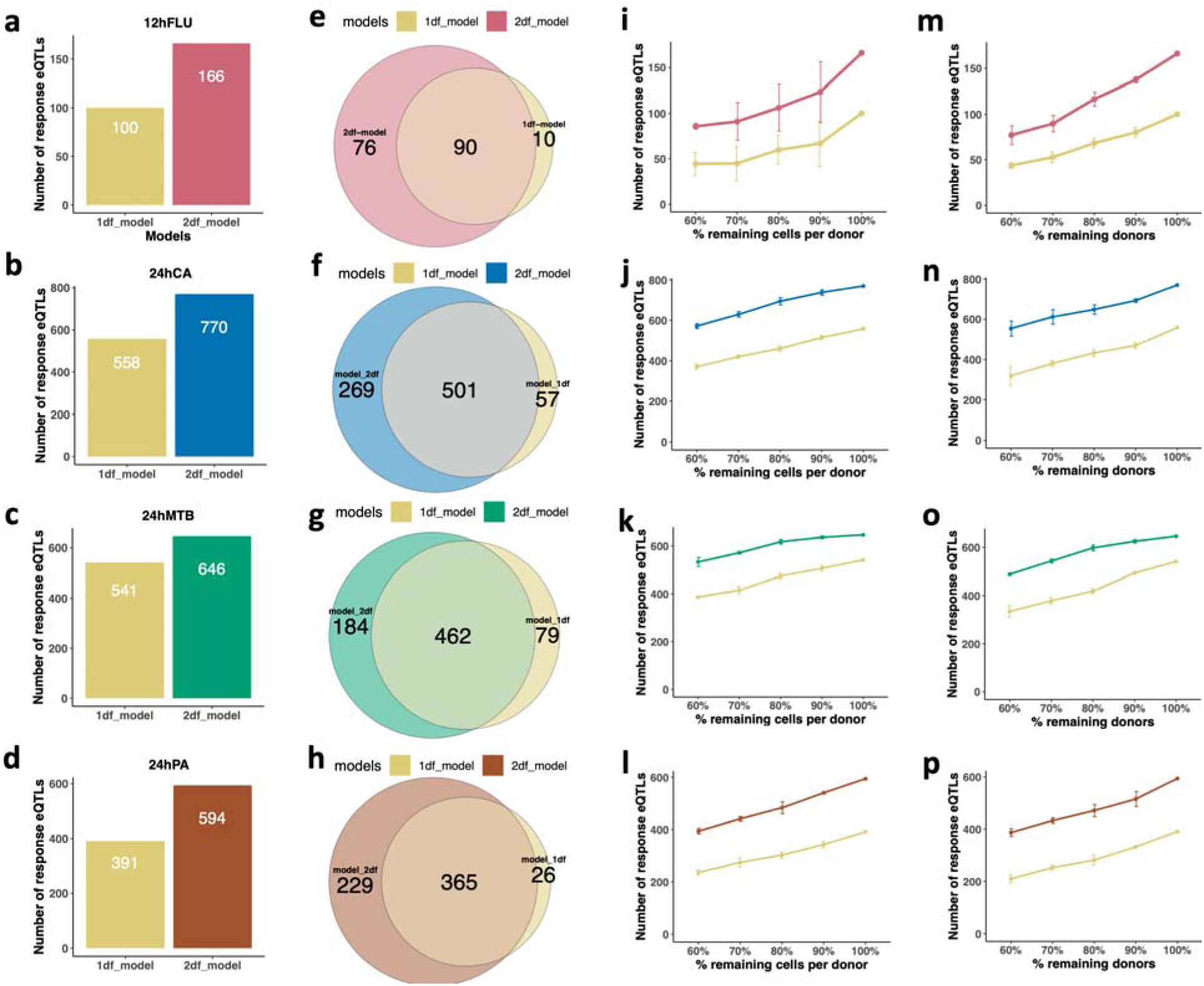
Mapping response eQTL in single cells under different perturbations. Modeling perturbation as discrete and continuous cell states (2df-model) found more response eQTL compared to an alternative model that considers only the interaction with the discrete perturbation state (1df-discrete) in IAV **(a)**, CA**(b)**, MTB**(c)**, and PA**(d)** perturbation experiments. Each bar represents the number of reQTL that were significant (Q value <0.05, P permutation<0.05) on each response eQTL model. **(e-h)** Venn diagram shows the overlapping of reQTL found by each model. the 2df-model identified on average 89.6% of the response eQTL discovered by the 1df-discrete across the 4 perturbations. In addition, on average 36.9% of the reQTL were found only by modeling the discrete and continuous perturbation state in the 2df-model. **(i-l)**. Plots show the number of reQTLs that remained significant (Q value<0.05) after randomly removing cells within each donors for each perturbation model. **(m-p)**. Plots show the number of reQTLs that remained significant (Q value<0.05) after randomly donors for each perturbation model.

**Figure 4.**
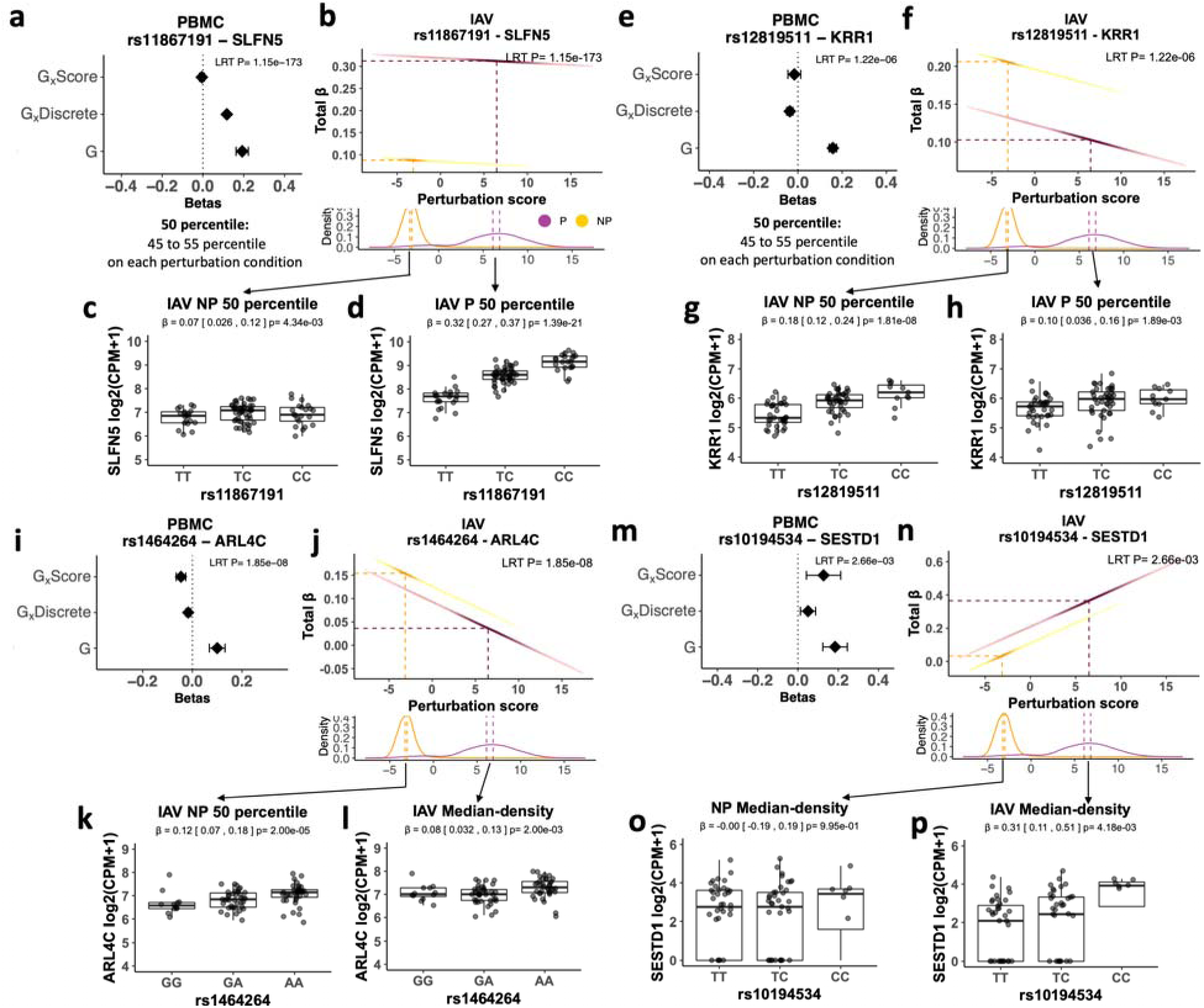
Estimating the change in the eQTL effect after perturbation in single cells. The 2df-model found substantial differences in the magnitude of the eQTL effect after IAV perturbation like the reQTL effect of rs11867191 for the gene *SLFN5* **(a-d)** and the reQTL rs12819511 for the gene *KRR1* **(e-h)**. Similarly, the 2df-model was able to detect subtle reQTL effects in the IAV experiment like the reQTL effect of rs1464264 for *ARL4C* **(i-l)** and the reQTL rs10194534 for *SESTD1* **(m-p)**. **(a, e, i, m)** Forest plots showing the effect size of each of the terms that contribute to the composite eQTL effect (total beta) for the eGene in the perturbation experiment. The effect sizes are from the PME 2df-model in PBMCs. **(b, f, j, n)** Scatter plots showing the total eQTL effect. Each dot represents the total beta in a cell. The color corresponds to the NP and P conditions. The dashed lines on the X axis indicates the perturbation score at the 50^th^ percentile. The dashed line on the Y-axis indicates the total beta at the 50th percentile. The histogram below the scatter plot shows the density of the perturbation score color by NP and P conditions. **(c-d, g-h, k-l, o-p)** Box plots showing the eQTL effect at the donor level. Each boxplot represents the eQTL effect around the 50 percentiles on the NP and P condition after aggregating the single cell gene count by donor (**METHODS**). G= genotype term, G_x_Discrete=Interaction term between G and perturbation, G_x_Score= Interaction term between G and the perturbation score.

Furthermore, when we focus on the previously prioritized eQTLs from Oelen et al 2022^21^, we observed an enrichment (**METHODS**) of significant interactions for CA (χ^2^ P=2.2e-178, **Figure S5 a**), PA (χ^2^ P=1.96e-137, **Figure S5 b**) and MTB (χ^2^ P=4.62e-143, **Figure S5 c**). These results demonstrated that our findings are driven by the G_x_E interaction effects.

**Figure 5.**
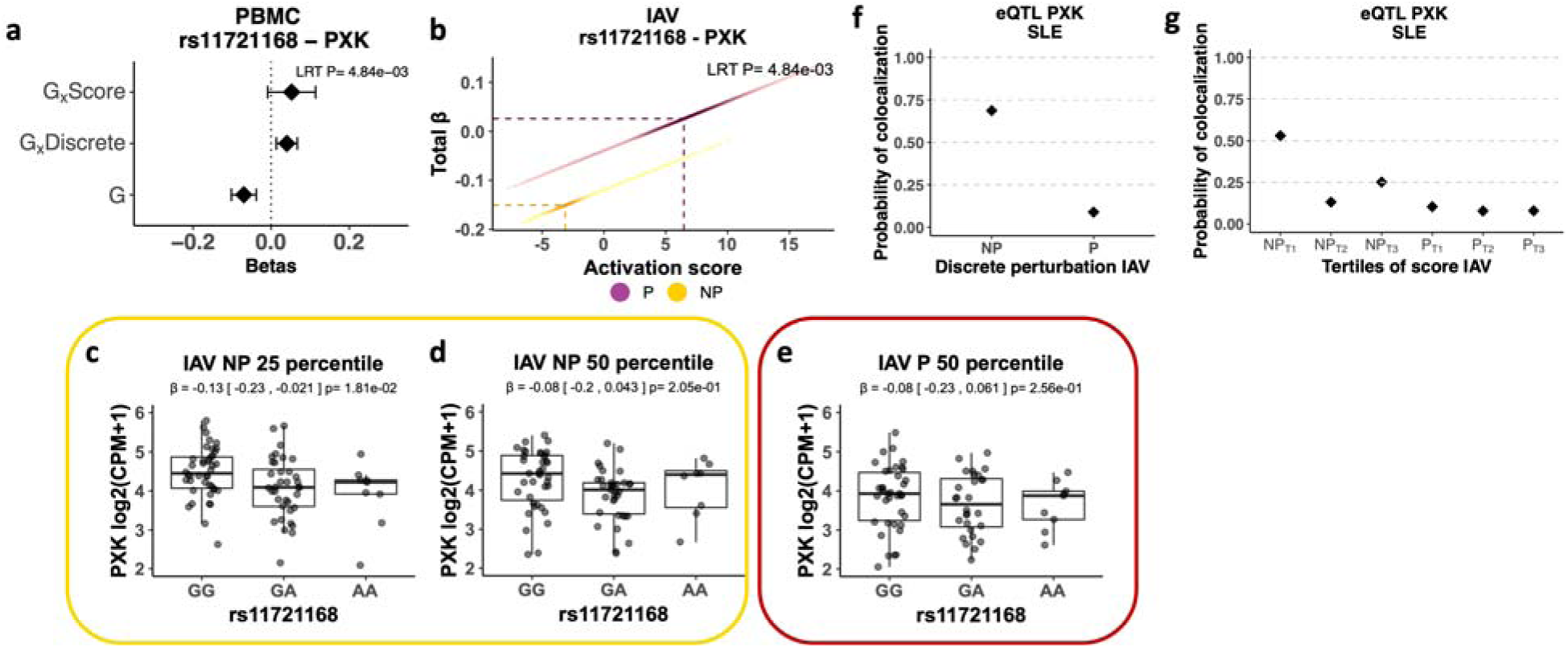
reQTL can have state-specific colocalization. We found that the reQTL rs11721168 for the gene *PXK* in the IAV perturbation experiment had perturbation-specific colocalization with systemic lupus erythematosus (SLE). **(a)**. Forrest plot shows the main genotype (G) and interaction effects (G_x_Score, G_x_Discrete) from the 2df-model in PBMC. **(b)**. Scatter plot showing the total eQTL effect of rs11721168 for *PXK* on each cell. The color corresponds to the NP and P conditions. The dashed lines on the X axis indicates the perturbation score at the 50th percentile. The dashed line on the Y-axis indicates the total beta at the 50th percentile. **(c)**. Boxplot represents the eQTL effect in cells between the 20th to 30th percentiles within the NP state after aggregating the single cell gene count by donor (**METHODS**). **(d-e)**. Boxplot represents the eQTL effect in cells between the 45th to 55th percentiles within each, the NP and P state after aggregating the single cell gene count by donor. **(f)**. Probability of colocalization (**METHODS**) of eQTL signal in *PXK* with GWAS for SLE at NP and P discrete state. **(g)**. Probability of colocalization of eQTL signal in *PXK* with GWAS for SLE at tertiles of the perturbation score (**METHODS**) in the NP (NP_T1_-NP_T3_) and P (P_T1_-P_T3_) cells. G= genotype term, G_x_Discrete=Interaction term between G and discrete perturbation state, G_x_Score= Interaction term between G and the continuous perturbation score.

### Comparison of the power to detect response eQTL effects

Our approach to discover reQTLs had two major features: (1) we use a single-cell model and (2) represent the perturbation state as a continuous variable. To test the effect of modeling single cells on power, we compared our 2df-model results to an approach that aggregates single-cell gene expression counts by condition and donor (Pseudobulk) and then tested the significance of the G_x_Discrete term using a linear mixed effects model (Pseudobulk-LME model) (**METHODS**). Across the 4 perturbation models the Pseduobulk-LME approach detected fewer reQTLs than the 2df-model (IAV: 26 reQTLs, CA: 238 reQTLs, PA: 106 reQTLs, and MTB: 484 reQTLs**, Figure S6 a-d, Table S1-S4**), and on average 72.8% of those were also found by the single-cell 2df-model (**Figure S6 e-h**). To address the effect of the continuous perturbation states on our power to detect reQTLs, we compared our 2df-model results to an alternative model that only considered the interaction effect of the genotype with the discrete perturbation state (G_x_Discrete) in a single-cell PME model (1df-discrete). We observed that including per-cell continuous perturbation state in the 2df-model still identified on average 89.6% of the reQTLs detected by the simple 1df-discrete model **(Figure 3 e-h)**; suggesting that little power was lost by including an additional term for continuous perturbation state interactions. Additionally, on average 36.9% of the reQTLs detected with the 2df-model were missed by the 1df-discrete model **(Figure 3 e-h)**.

**Figure 6.**
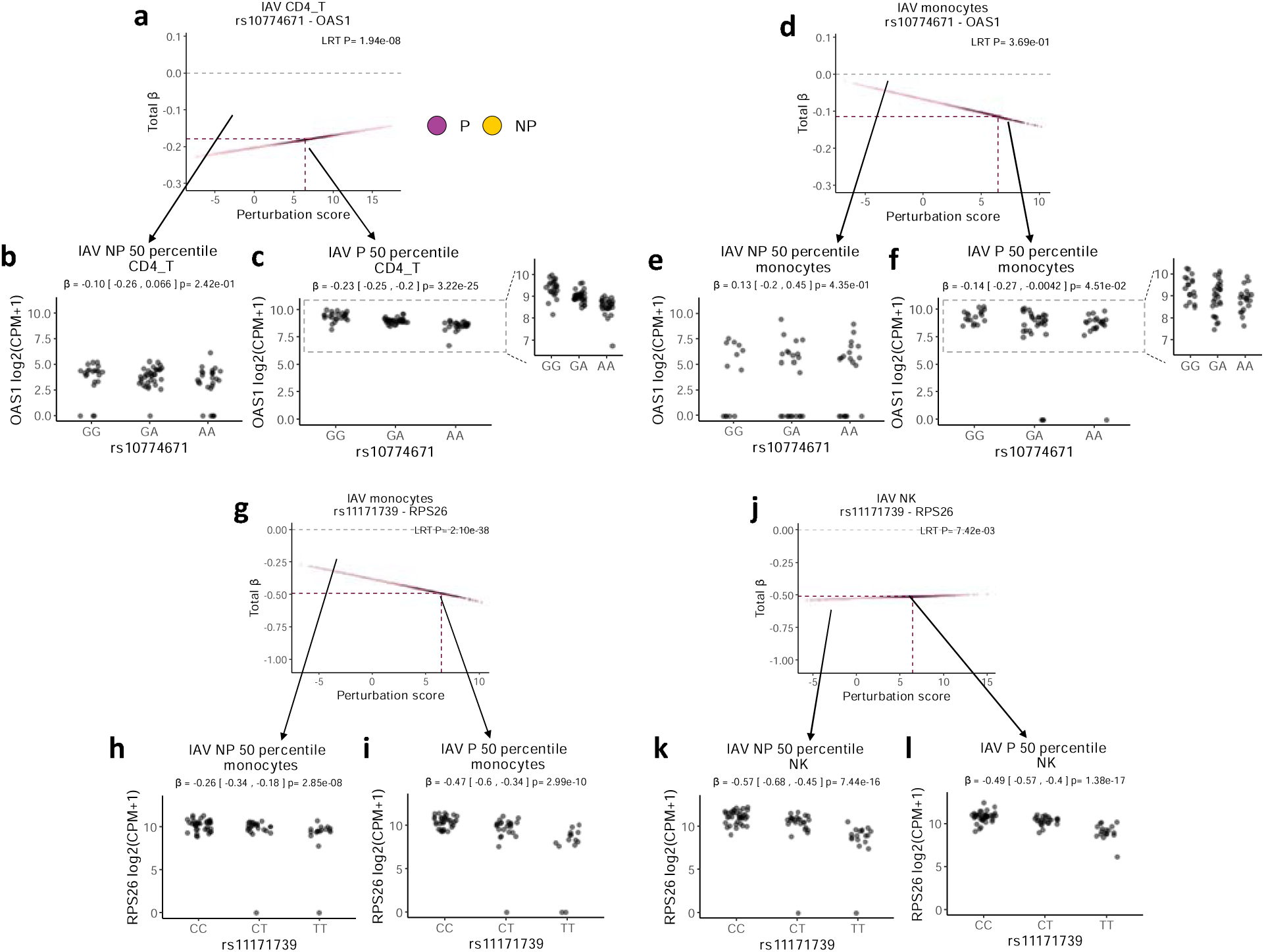
Response eQTL with cell-type effects. We observed that an average of 25% of the reQTL have heterogeneous effects across cell types. For example, we found that in the IAV experiment the eQTL rs10774671 for the gene *OAS1* was magnified after perturbation in all cell types, however the magnitude of the effect was stronger in CD4-T cells **(a-c)** but more subtle in other cell types like monocytes **(d-f).** Furthermore, we found that the reQTL rs1117139 for the gene *RPS26* had cell-type specific effects. After perturbation, the eQTL effect was magnified in monocytes **(g-i)**, however the eQTL effect decreased in NK cells **(j-l).** Scatter plot showing the total eQTL effect. Each dot represents the total beta in a cell. The color corresponds to the NP and P conditions. The dashed lines on the X axis indicates the perturbation score at the 50th percentile. The dashed line on the Y-axis indicates the total beta at the 50th percentile. Each boxplot represents the eQTL effect around the 50 percentile on the NP and P condition after aggregating the single-cell gene count by donor (**METHODS**). **(c,f)** the area delimited by the dashed box was magnified.

Next, we tested if these differences in power remained consistent under different sample size scenarios. We compared the effect on the power to detect reQTLs by randomly reducing the number of cells per donor (**Figure 4 i-l**) and also randomly reducing the number of donors (**Figure 4 m-p**). We compared the power difference between the 2df-model and the simplified 1df-discrete. We found that the 2df-model consistently found more reQTLs under any of the evaluated scenarios. Also, we observed a linear power loss by reducing donors and cells, suggesting that future studies with larger sample sizes may be able to detect novel response eQTL effects using the 2df-model.

### reQTL with strong and subtle perturbation effects

Our model allows us to define a total eQTL effect for each cell. This effect is the sum of the main genotype effect (G) and the interaction effects (G_x_Discrete and G_x_Score) multiplied by their corresponding perturbation state for each cell (**METHODS**). We observed eQTLs with substantial change in the magnitude of effect upon perturbation. For example, the effect of each of the interaction terms and the main genotype effect of rs11867191 on *SLFN5* (G_PBMC_=0.28, P=4.82e-36, G_x_Discrete_PBMC_=0.11, P=4.6e-174, G_x_Score_PBMC_=-0.005, P=3.8e-01, **Figure 4a**). This resulted in a substantial change in the total eQTL effect of rs11867191 on *SLFN5* after perturbation (**Figure 4b**, 2-df model_PBMC_ LRT P=1.15e-173, **Table S2**). To confirm the change in the *SLFN5* eQTL at the donor level, we selected cells between the 45th and 55th percentile (50p) of the perturbation score distribution from within perturbed (P) and non-perturbed (NP) state (**METHODS**). We found that the eQTL effect in the NP state (NP-50p eQTL_PBMC_=-0.07, LRT P=4.34e-03, **Figure 4 c**) was magnified upon perturbation (P-50p eQTL_PBMC_=-0.32, LRT P=1.39e-21, **Figure 4 d**). A similar example, that replicated across the four perturbation models was the reQTL for *KRR1* (IAV: rs12819511 2-df-model_PBMC_ LRT P=1.22e-06, CA: rs35619460 2-df-model_PBMC_ LRT P=2.37e-38, MTB: rs35619460 2-df-model_PBMC_ LRT P=1.82e-32, and PA: rs35619460 2-df-model_PBMC_ LRT P=5.49e-27, **Table S2-5**, **Figure S7 a-i**). In the IAV experiment, the reQTL has a strong interaction effect with the discrete perturbation state and an additional independent interaction with the continuous perturbation state (**Figure 4e-h**).

In addition, our model was able to detect more subtle changes in reQTL effects, such as the reQTL rs1464264 for the gene *ARL4C* (IAV 2-df-model_PBMC_ LRT P=1.85e-08, **Table S2**), where we observed the independent contribution of the two interaction effects (G_x_Discrete_PBMC_=-0.01, P=9.5e-04; G_x_Score_PBMC_=-0.04, P=3.6e-06, **Figure 4 i**) on the attenuation of the total eQTL effect in *ARL4C* (**Figure 4 j**). This reQTL was still significant in the reduced single cell model (1df-discrete_PBMC_ LRT P=1.11e-04, **Table S2**), however it was not detected by the Pseudobulk-LME model (Pseudobulk-LME_PBMC_ LRT P=2.07e-01, **Table S2**). A similar example was found in the reQTL rs10194534 for the gene *SESTD1*, where the G_x_Score_PBMC_ term was the most significant interaction effect in the 2df-model (LRT P= 2,66e-03, **Figure 4 m**), and this reQTL was not significant in the model that only include the discrete perturbation effect (1df-discrete_PBMC_ LRT P=6.36e-02, **Table S2**). We confirmed these results by looking at the eQTL effect of these genes in the NP-50p and P-50p state (*ARLC4* **Figure 4 k-l** and *SESTD1* **Figure 4 o-p**). Our findings in *SLFN5*, *KRR1 ARL4C*, and *SESTD1* are consistent with previous studies that identified reQTL effects in perturbations experiments that induced an interferon response^10,11^.

### reQTLs identify genes with perturbation-specific regulation

Finding eQTLs that shared their causal variant with a trait (colocalization) can be limited, in part because the eQTLs effects are typically not ascertained in the relevant context, e.g., perturbed states, thus reQTLs may be more likely to colocalize compared to an eQTL that has no change upon perturbation. We performed a colocalization analysis with 9 immune and 4 non-immune traits (coloc.abf, probability of colocalization Pr>50%. **METHODS**). As expected, we found that reQTLs identified with the 2df-model in all perturbation experiment except for MTB were enriched (**METHODS**) for colocalizations (IAV P=1.03e-04, CA P=2.73e-04, MTB P=2.24e-01, and PA 8.91e-06 and **Figure S8**). Furthermore, 558 of 2,176 reQTLs detected in the 2df-model were not detected by the 1df-discrete model. For example, in the CA experiment, rs3184122 was a reQTL for the gene *COX14* (**Figure S9 a-c**), and has also been associated with lymphocyte counts^27^, and reQTL rs2238641 for the gene *FARSA* (**Figure S9 d-f**) was previously associated with hematocrit levels^28^. In the PA experiment we found the reQTL rs3108680 for the gene *PRMT6* (**Figure S9 g-i**), was previously associated with levels of low-density lipoprotein cholesterol (LDL)^29^.

In some cases, the reQTLs detected only in the 2df-model pointed to perturbation-specific disease relevance. One such example was the reQTL effect of rs11721168 for *PXK* in the IAV experiment (IAV 2df-modelPBMC LRT P=4.84e-3, G_PBMC_=-0.10, P=1.74e-05, G_x_Discrete_PBMC_=0.06, P=3.38e-03x; G_x_Score_PBMC_=0.08, P=9.30e-02, **Table S2** and **Figure 5 a**). We observed that there was a decrease in the eQTL effect of rs11721168 for *PXK* after perturbation (**Figure 5 b**). We confirmed that the eQTL effect was presented in cells that score lower in the non-perturbed (NP) state (**Figure 5 c**), and the eQTL was no longer significant at either the 50p of the NP state (**Figure 5 d**) or the 50p of the perturbed state (**Figure 5 d**). In addition, we found that the eQTL signal for *PXK* in the NP cells colocalized with a GWAS locus for SLE^30^ (**Figure 5 f**). When we performed the colocalization at increasing tertiles of the perturbation score within the NP-state (NP_T1_-NP_T3_) and P-state (P_T1_-P_T3_) (**METHODS**), we found that the eQTL signal for *PXK* colocalized in the lowest tertile of the NP cells (colocalization Pr _NPT1_=53.1% **Figure 5 f**). *PXK* encodes a sorting nexin (SNX) protein important for receptor internalization, organelle trafficking including endosomal trafficking^31^, and it has been previously implicated in the risk of SLE^32,33^. Our results shows that genetic regulation of *PXK* is particularly strong in lowly perturbed cell states. It also suggests that these may be the most relevant cell states through which *PXK* may be mediating the risk for SLE. A similar example was observed with the reQTL rs3807865 for *TMEM106B* (**Figure S10 a-d**) that showed perturbation-specific colocalization with coronary artery disease (**Table S11-S12**) and depression (**Figure S10 e-f**).

### Response eQTLs can have cell-type-specific effects

Previous studies demonstrated that eQTLs can have cell-type dependent effects^34^. Because our response eQTL analysis was performed in samples of heterogeneous cell-type composition, we explored the possibility of heterogeneous reQTL effects between cell-types. To evaluate this, we fitted the single cell 2df-model for each cell type and then we tested the heterogeneity of each of the interaction terms between cell types using the Cochran’s Q-test (QE, **METHODS**). We consider reQTLs with heterogeneous effects as those where at least one of the interaction terms (G_x_Discrete or G_x_Score) have a QE test Q value<0.10. We found that most reQTLs have similar effects across cell types, however around 25% of reQTLs showed evidence of heterogenous response eQTL effects across the 4 perturbation models (**Figure S11**. 40/166 for IAV, 202/770 for CA, 209/594 for PA, and 145/646 for MTB, **Table S10**).

For example, we observed that the reQTL rs11867191 for *SLFN5* had cell-type specific effects in the IAV experiment, while the interaction effects between the genotype and the perturbation state (G_x_Discrete) were not significantly different (QE_GxDiscrete_ P=0.25) across cell types (G_x_Discrete_B_=0.13, 95%CI: 0.08,0.18; G_x_Discrete_CD4-T_=0.11 [95%CI: 0.10,0.12]; G_x_Discrete_CD8-T_=0.08 [95%CI: 0.06,0.10]; G_x_Discrete_Monocytes_=0.09 [95%CI: 0.06,0.12]; G_x_Discrete_NK_=0.06 [95%CI: 0.03,0.08]. **Table S6**), the interaction effects between the genotype and the continuous perturbation state (G_x_Score) showed evidence of heterogeneity (QE_GxScore_ P=3.06e-4) between cell types (G_x_Score_B_=-0.04 [95%CI: -0.10, 0.007]; G_x_Score_CD4-T_=0.01 [95%CI: 0.001, 0.03]; G_x_Score_CD8-T_=0.004 [95%CI: -0.03, 0.04]; G_x_Score_Monocytes_=0.02 [95%CI: -0.04, 0.09]; G_x_Score_NK_=0.02 [95%CI: -0.03, 0.07], **Table S6**). As a result, we observed that the total eQTL effect in B cells (**Figure S12 a)** and monocytes (**Figure S12 d**) was different than in other cell types (**Figure S12 g, j, m**). Confirming this, the eQTL effect in the P-50p in B cells (eQTL_B_=0.40 [95%CI: 0.31,0.49], **Figure S12 c**) and monocytes (eQTL_monocyte_=0.41 [95%CI: 0.26,0.55], Figure S12 f) was stronger compared to the eQTL in the P-50p in CD4-T (eQTL_CD4-T_=0.29 [95%CI: 0.24,0.34], **Figure S12 l**), and the other cell types (**Figure S12**).

Similarly, we found that the reQTL rs10774671 for the gene *OAS1* in IAV (2df-model_PBMC_ LRT P=3.84e-15) had cell-type-specific interaction effects (QE_Perturbation_ P=6.7e-03, QE_GxScore_ P=7.2e-02). We found that the increase in the total eQTL effect was more pronounced in CD4+ T cells (**Figure 6 a**) and B cells (**Figure S13 a**) while the increase in the total eQTL effect was smaller in monocytes (**Figure 6 d**), CD8+ T (**Figure S13 d-f**) and NK cells (**Figure S13 g-i**).These differences were also observed at the donor level, where we found that the eQTL effect in the P-50p of CD4+ T cells (β_CD4-T_=-0.23 [95%CI: -0.25,0.20], **Figure 6 b-c**) and B cells (β_B_=-0.20 [95%CI: -0.25,-0.15], **Figure S13 b-c**) were stronger than the eQTL effect at P-50p in monocytes (β_Monocyte_=-0.14 [95%CI: -0.27,-0.004], **Figure 6 e-f**). Furthermore, we found that the eQTL rs10774671 for *OAS1* had state-specific colocalization with the GWAS locus for severe COVID-19 (Pr_NP_=6.7%, Pr_P_=98.6%, **Figure S14 a** and **Table S11**). Similarly, we observed colocalization within tertiles of the perturbation score in the perturbed cells (Pr_PT1_=95.7%, Pr_PT2_=98.7%, Pr_PT3_=98.7%, **Figure S14 b** and **Table S12**). The relationship between the genetic variant rs10774671, the levels of *OAS1* and the risk of severe COVID19^35^ (rs10774671-A, OR=1.10, P=6.0e-10) has been previously reported. Briefly, the *OAS1* gene encodes an interferon stimulated protein that activates Ribonuclease L which then degrades cellular and viral RNA^36^. Furthermore, lower levels of this protein contributed to the severity of COVID-19^37^. Our result suggests that cell types like B and CD4+ T cells may have a more substantial contribution to the interindividual differences in the levels of *OAS1* in the context of cell perturbation by an RNA virus.

We also observed the reQTL rs11171739 for the gene *RPS26* in the IAV experiment (2df-model_PBMC_ LRT P=4.24e-165, **Table S2**). This reQTL has cell-type-specific interaction effects (QE_GxDiscrete_ P=1.73e-29, QE_GxScore_ P=2.14e-54). More specifically, we found that there was a significant magnification in the total eQTL effect after perturbation in monocytes (**Figure 6 g**) and B cells (**Figure S13 j-l**), however the change in the eQTL effect was not only smaller in NK cells (**Figure 6 j**) and CD4+ T (**Figure S13 m-o**) but strikingly, also in the opposite direction. Furthermore, there was no significant reQTL effect in CD8+ T cells (**Figure S13 p-r**). These differences in gene regulation in the context of perturbation were also observed when we estimated the eQTL effect on NP-50p and P-50p states on monocytes (**Figure 6 h-i**), NK cells (**Figure 6 k-l**) and the other cell types (**Figure S13**). We observed similar cell-type-specific reQTL effects in the *RPS26* gene in the CA, PA and MTB perturbation experiments (**Figure S15-S17**). In addition, we found that the eQTL signal for *RPS26* in IAV colocalized with a GWAS locus for RA^38^ in both discrete perturbation states (Pr_NP_=88.9%, Pr_P_=87.7%, **Figure S14 c** and **Table S11**), and we did not observed differences in the strength of colocalization along the tertiles of the perturbation score (**Figure S14 d, Table S12**). Previous studies suggest variant rs11171739 has pleiotropic effects on autoimmune diseases^39^ and increases the risk for RA^38^. Our results show that genetic regulation of *RPS26* depends on cell perturbation and cell-type, both of which may influence the causal relationship with diseases like RA.

## Discussion

Recent studies have extended the analysis of reQTLs to single-cell data. However, the identification of reQTLs using linear models has been limited to the comparison of effects between the discrete non-perturbed and perturbed experimental conditions. In addition, the analysis is typically stratified to the cell-type level, further reducing the power to detect reQTL effects. Here, we introduced a framework that incorporates the level of at which each cell responds to the perturbation to model reQTL effects across different continuous perturbation states like interferon alpha, complement activation and inflammatory response. We demonstrated a substantial increase in power to detect reQTL effects compared to models that ignore cell-level variation in the response state of a cell. In addition, we demonstrated that our framework can be applied to heterogeneous samples like PBMCs and further examined the differences in reQTL effects between cell types.

The limited colocalization between eQTLs and GWAS loci suggested that disease alleles may affect gene regulation in specific cellular contexts, which may be difficult to capture on aggregated eQTL profiles. We observed an enrichment of reQTLs’ colocalization with traits, which highlights the importance of improving the identification of reQTLs. Consistent with previous studies^8,11^, we found that reQTLs can have perturbation-dependent colocalization. In addition, we found that even among cells with the same discrete perturbation state, the strength of colocalization of the reQTL and a trait can depend on the continuous perturbation state of the cell. This suggests that in a reQTL study, identifying the most responsive cell state may offer the opportunity to expand the understanding of the genetic mechanism of disease.

Our reQTL model has important limitations. First, our PCA-based perturbation scores define the continuous cell states within each perturbation dataset, which may not transfer to other independent datasets, even with the same perturbation condition.

Second, we did not define the perturbation score independently for each cell type, because the perturbations lasted nor more than 24 hours, which can induce early changes in gene regulation, but may not reach later-stage differentiation states such effector cytotoxic CD8+ T or Th-1 CD4+ T cells^40–43^. Thus, *ex-vivo* perturbation studies that seek to capture cell-type-specific changes in more differentiated cells may require the development of cell-type specific scores. Third, the perturbation score presented here is based on a log-linear model, but scores that aim to capture cell-type specific programs may require a different approach like non-linear models of perturbation.

Overall, however, our work provides a general framework for better detection of perturbation-dependent eQTLs and underscores the importance of modeling cell-level variation. Combining these statistical models with biologically relevant perturbations will ultimately advance our understanding of the biologic mechanisms that underlie disease heritability.

## Acknowledgements

This work is supported in part by funding from the National Institutes of Health (R01AR063759, P01AI148102, U01HG012009, UC2AR081023, R56HG013083), T32DK110919 and T32HG10464-5 (C.V), F30AI172238 (J.B.K.), T32GM007753 and T32GM144273 (J.B.K., L.R.)

## Code availability

A GitHub repository containing the scripts for reproducing the analyses in the manuscript will become available upon acceptance.

## Methods

### Cohorts

We used published data (GEO accession number GSE162632) to study the effect of 12 hours of *ex-vivo* stimulation with either influenza A virus (IAV) or mock (control) in 90 samples of heterogenous cell composition (peripheral blood mononuclear cells, PBMCs) using single cell RNA-seq. Similarly, we obtained data (EGA accession number EGAS00001005376) of 120 PBMCs samples infected for 24 hours with *Candida albicans* (CA), *Pseudomonas aeruginosa* (PA), *Mycobacterium tuberculosis* (MTB) and mock (control) from the European Genome-Phenome Archive under a Lifelines DEEP DAG2+ Project (“LLDEEP DAG2+”) Data Access Agreement.

### Genotype quality control

The original Randolph, et al. study (IAV perturbation) used 4x low-pass whole-genome sequencing to call genotypes of 89 individuals a total of 78,111,311 genome-wide variants. Genotypes with a posterior probability (GP) <0.9 were considering as missing data. We removed variants with a missing call rate>10% and only included variants with a minor allele frequency >=5% and Hardy-Weinberg equilibrium (HWE) P>1e-05. In total, for the IAV perturbation dataset we had 5,194,816 genetic variants genomewide. Furthermore, we selected 508,883 independent variants using plink^44^ (-- indep-pairwise 50 5 0.2) and then we performed principal component analysis using the R package SNPRelate^45^.

The original Oelen, et al. study (CA, PA and MTB perturbations) used genotype data from the HumanCytoSNP-12 BeadChip and imputed with the Haplotype Reference Consortium (HRC) reference panel of 120 individuals at a total of 7,249,882 genomewide variants mapped on the Hg19 reference genome. For the main analysis we included variants with missing call rate <10%, minor alle frequency (MAF)>=5% and Hardy-Weinberg equilibrium (HWE) P>1e-05. The variants that passed the filtering step were then lifted over to the hg38 reference genome using Picard tools. In total, for this dataset, we had 3,962,615 variants. We selected 292,681 independent variants using plink^44^ (--indep-pairwise 50 5 0.2) and then we performed principal component analysis using SNPRelate^45^. For the secondary analysis, we selected the 3,554 variants included in the Supplementary table 9 of Oelen et al and successfully lifted over 2,347 variants from the hg19 to hg38 reference genome.

### Single cell mRNA quality control

We performed quality control of the 236,993 PBMCs from the 12 hours perturbation experiment with IAV. These cells had <20% reads that mapped to mitochondrial genes. We removed 10 influenza IAV genes from the original 19,248 genes in the count matrix. We selected a total of 208,223 cells with >500 expressed genes for the analysis. For the 24-hour perturbations with CA, PA and MTB there were, in total, 493,061 cells and 33,538 genes measured. As in the IAV perturbation study, we selected 475,333 cells with <20% reads mapping to mitochondrial genes and >500 detected genes.

Then, for each perturbation condition (NP or P) in each study, we normalized the counts for each cell, scaled each gene’s expression, and cosine normalized. We selected the 3000 most variable genes based on the variance stabilizing transformation (VST) and conducted principal components analysis (PCA) using an implicitly restarted Lanczos method implemented in the irlba R package. Then, we controlled the effects of batch and sample donor using Harmony^46^ (V0.1.0) (parameters: θ_donor_ = 1, θ_batch_ = 1). The top 20 corrected expression PCs (hPCs) were used for downstream analysis. For cell-type annotations, we used the originally published cluster labels.

### Estimating the continuous perturbation state of a cell

For each perturbation model, we developed a continuous perturbation score of a cell by modeling the discrete perturbation state (Non-perturbed=0, Perturbed=1) as a function of the top 20 hPCs in a logistic Lasso regression model implemented in the R package glmnet^47^. We fitted the model and used the LASSO penalty term (λ) with a tuning parameter obtained from cross-validation (cv.glmnet parameters: alpha=1, lambda.1se)

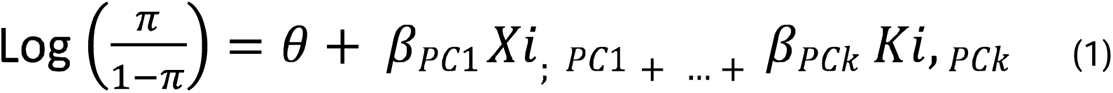

Where K is the number of PCs and π is the probability of the cell to be in the pool of perturbed cells.

### Gene pathway enrichment analysis

For each perturbation experiment, we calculated the Pearson correlation between the 3000 most variable genes and the perturbation score of a cell. We selected genes with a correlation Q value <0.05. Then, we use these correlation values to rank the genes and performed a gene pathway enrichment analysis using the R’s fgsea package^25^ with a maximum gene set size of 500, a minimum size of 15, and 100,000 permutations. We include the following gene sets from the Pathways Interaction Database (“PID”) stored in MSigDB’s: Hallmark gene set and large collection of immunological conditions (C7).

### Prioritizing eQTLs in PBMCs bulk expression profiles

For each individual and perturbation-condition we summed the gene expression counts for each gene across all cells to generate bulk profiles, and then, we averaged the gene expression from each perturbation condition by individual resulting in the average bulk expression profiles. For each perturbation experiment we generated bulk expression profiles for the NP, P and average conditions. We normalized the expression profiles (log2 CPM+1) and inverse variance transformed. Then, we removed the effect of latent factors using PEER factor analysis^48^. We only included transcripts expressed in at least 50% of the samples. The corrected gene expression was used as the dependent variable in our bulk cis-eQTL mapping. We performed the cis-eQTL mapping by testing each gene with genetic variants within 1 MB of each gene’s transcription starting site (TSS) using a linear regression implemented in FastQTL^49^. We use 1000 permutation per locus and select the gene-SNP pairs (eGenes) with a Q value<0.05. In most cases, we prioritized the eQTL from the averaged NP and P profile as this produced the largest number of eQTLs (Q value<0.10) (**Figure S3**).

### Identifying response eQTL effects

For the single cell approaches, we modeled the raw expression (E) counts in single cells as a function of genotype allele dosage and an interaction with cell state accounting for confounders and technical batch using the Poisson Mixed Effects model (PME) described by Nathan et al 2022, Nature^17^. For all the analyses, we included age and sex as a fixed effect, where available. We also controlled for genetic population stratification by including the top 5 genotype PCs (gPC) as covariates. Similarly, we controlled for unwanted variation in gene expression by including the top 5 gene expression PCs (ePC). In addition, we included the number of unique molecular identifier (nUMI) and percentage of reads mapping to mitochondrial genes (MT%) as fixed effects, and donor (d) and technical batch (b) as random effects. We include two cell states variables, the discrete perturbation state (Discrete: where P=1 for perturbation or 0 for no perturbation) and the residual perturbation score (rScore) (3) that avoid the collinearity between the Discrete variable and the perturbation score (Score) from equation (1). To estimate the reQTL effect, we fit a model that included the genotype main effect (G) and the interaction terms of G with the Discrete (G_x_Discrete) and rScore (G_x_rScore) variables. In contrast, we only included the interaction term G_x_Discrete in the simplified 1df-discrete model.

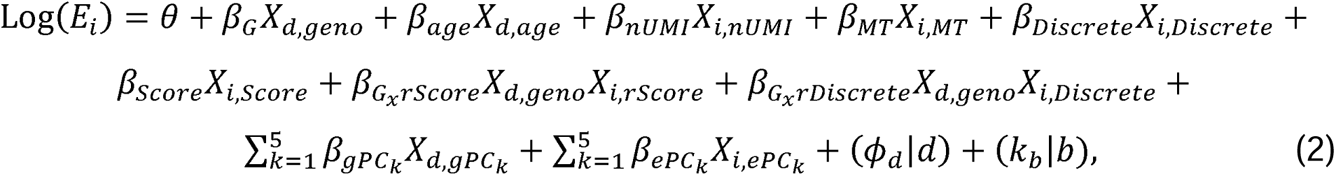

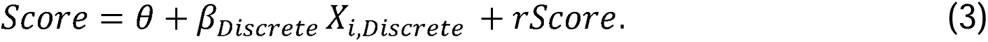

To map reQTLs in bulk profiles (Pseudobulk-LME) we modeled the residuals from the PEER factor analysis (rPFA) as a function of the genotype and the G_x_Discrete term accounting for donor’s age, sex, and both the total numbers of UMIs and mean MT% across all cells per donor using a linear mix effect (LME) model. We included donor as a random effect in the LME model (4).

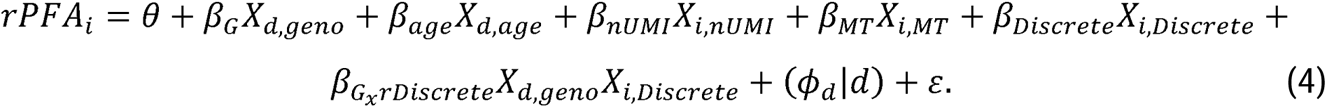

We compared the significance of each of these full models against a null model with no interactions terms using the likelihood ratio test and considered significant reQTL to be those with a LRT Q-value<0.05. Furthermore, to rule out the possibility of spurious significant results, we performed 500 permutations per each discovered reQTL. For the permutation in the single cell models, we randomly shuffled values of the perturbation score and the discrete perturbation state within each donor. For the pseudobulk-LME model, we randomly shuffled values of the perturbation state across samples.

To exclude spurious associations, for each gene we counted the proportion of LRT-P values from the permutation experiments that were smaller than the discovery LRT-P (permutation P) and removed genes with permutation P <0.05. To exclude genes where the PME model may not be appropriated due to high differential gene expression across cell states, we fit a PME model where we simulated null conditions by randomly shuffling the perturbation score value within each donor. We tested the significance of the interaction term under this null by comparing this model with a model with no GxScore term using the Likelihood ratio test (LRT ANOVA). For each eGene we performed 100 permutations and removed eGenes where we observed a significant P value (LRT P<0.01) in more than 5 out of 100 permutations.

In addition, we performed a secondary analysis for the CA, PA and MTB perturbation experiments based on the 1825 eGenes prioritized by Oelen et al and listed on supplementary table S9 and passing QC in our pre-processing. For each perturbation experiment we estimated reQTL effects for this set of 1825 eQTLs using each of the models described above (equations 1 and 3) and compared the significance against a null model with no interactions terms using the LRT and considered significant those with LRT Q-value<0.05. Furthermore, for each perturbation experiment we generated p values under the uniform distribution based on the number of eQTLs included in our secondary analysis. Then, we calculated the enrichment of significant reQTLs in the 2df-model by comparing the number of test where we found 2df-model LRT P< 0.001, >0.001 & <=0.01, >0.01 & <=0.05, >0.05 & <=0.1, and >0.1 with those found under the uniform distribution using a χ2 test.

### Estimating the eQTL effect at levels of the perturbation score

For any given reQTL, we selected cells either in the NP or P condition and calculate the 45^th^ and 55^th^ percentile of the perturbation score in that group. Then, we select the cells with perturbation score values within the 45^th^ to 55^th^ percentile (50p) and estimated the eQTL effect in those cells by fitting a single cell PME, where we modeled the raw gene counts as a function of the genotype (G) and the same cell and donor level fix and random covariates as described above (equation 2) but without interaction terms (5). We consider the significance of the G term by comparing with a null model with no G term using the LRT with 1 degree of freedom.

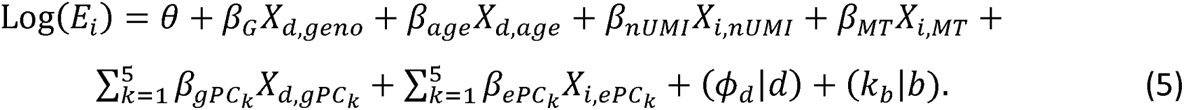

To visualize the eQTL effect at the donor level, we aggregated the gene counts in the cells in the 50p by donor and then normalized the gene expression (log2 CPM+1). Then, we plot the gene expression as a function of the eQTL’s allele dosage.

### Composite eQTL effect in a cell

We defined the total eQTL in a cell as the sum of the genotype main effect with each of the interaction effects (G_x_Discrete) and G_x_Score, weighted by its corresponding value (Discrete: NP=0, P=1,Score: continuous perturbation score) (6).

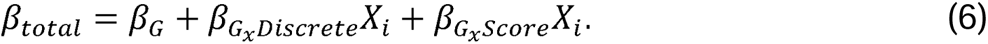

### Comparison of Poisson mixed effect and negative binomial models

To confirmed that the single cell PME model was well calibrated, we fit a single cell reQTL model where we modeled the raw expression counts in single cells as a function of genotype allele dosage and an interaction terms equivalent to the PME 2df- model described in equation 1, using a negative binomial model implemented in the glmmTMB^50^ R package. We calculated the correlation between the LRT-P from the original PME and negative binomial model.

### Down-sampling donors and cells

To test the effect of sample size on the power to detect reQTLs in the single cell models, we generated data sets where we randomly removed cells or removed donors. We generated 5 data sets, and in each, we randomly removed either 10, 20, 30, 40, or 50% of cells within each donor. For each data set we generated 3 replicates by using different random seeds. Similarly, we generated 5 data sets with three replicates each where for each set we randomly removed either 10, 20, 30, 40, or 50% of donors. Then, in each downsampled data set, we estimated reQTL effects in with the PME model by testing the originally prioritized eGenes and counting the number of reQTLs that reached a LRT Q-value<0.05.

### Response eQTL with heterogeneous effects across cell types

We tested for heterogeneous effects by comparing the effect size of each of interaction terms between cell types. For each perturbation experiment, we selected cells from each major cell type presented in the experiment (IAV: B, CD4+ T, CD8+ T, Monocytes, NK; CA-PA-MTB: B, CD4+ T, CD8+ T, Monocytes, NK, DC) and fitted the single cell 2df-model with the G_x_Discrete and G_x_Score terms. Then, for each reQTL we use the summary statistics to compare the effect sizes of each of the interaction’s terms across cell-types. We tested if the variability in the observed effect sizes is larger than the expected based on sampling variability alone using the Cochrane’s Q-test (Q) implemented in the metafor R package (rma, method=’FE’, weighted=TRUE). We defined response eQTL with heterogeneous effects as those with a Cochrane’s Q-test Q-value<0.10 in at least 1 of the interaction terms.

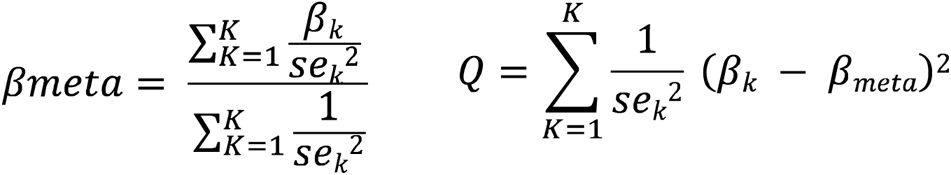

Where, K is the number of cell-types in the perturbation experiment, βk is the effect size for each cell-type, and β_meta_ is the average effect size weighed by the inverse of the variance of the estimate on each cell-type.

### Colocalization analysis

To find instances where a perturbation-dependent reQTL shared its causal variant with a GWAS locus, we performed a colocalization analysis using coloc^51^ (coloc abf), a Bayesian approach that estimates the posterior probability of having a shared true causal variant between the eQTL and the GWAS locus. We used summary statistics of 9 GWAS on immune traits (Type 1 diabetes^52^,Multiple sclerosis^53^, Asthma^54^, Rheumatoid arthritis^38^, systemic lupus erythematosus^30^, inflammatory bowel disease^55^, Crohn’s disease^55^, ulcerative colitis^55^, psoriasis^56^) and 3 GWAS on non-immune traits (coronary artery disease^57^, type 2 diabetes^58^, depression^59^).

We defined eQTL as those with a nominal Q value(P)<0.05 in the analysis with FasQTL. We ran coloc with the default parameters and defined colocalization as those loci with a posterior probability H_4_>0.5. Furthermore, for those colocalized loci, we generated the 95% credible set and obtained the posterior probability of each variant withing the credible set (snpH_4_) under the single causal variant assumption. We calculated the enrichment of colocalization in eGenes with reQTL effect compared to eGenes with no reQTL effect using Fisher’s exact test.

In addition, to find colocalization at different levels of the perturbation score, we defined tertiles of perturbation within the NP (T1-T3) and P (T4-T6) conditions. For example, in the IAV experiment, T1 was defined as NP cells with perturbation score values <33^th^ percentile (-0.90), T2 was defined as NP cells with the perturbation score >=33^th^ and below 66^th^ percentile (-0.75), and T3 was defined as NP cells with an perturbation score >= 66^th^ percentile. Then, for each tertile we generated pseudobulk gene expression profiles, tested for cis-eQTL and ran coloc as described before.

## Supplementary information

### Supplementary Tables

**Table S1. Gene set enrichment analysis**. (See Valencia_etal_TableS1_GSEA.xlsx)

Gene set enriched for each of the perturbation score. MSigDB gene set (GSEA P adjusted<0.05)

**Table S2. Results from reQTL analysis in IAV.** (See Valencia_etal_TableS2_IAV_reQTL_Models.xlsx)

Results from the 2df-model (tab 1), 1df-discrete (tab 2) and pseudobulk LME model (tab 3).

**Table S3. Results from reQTL analysis in CA.** (See Valencia_etal_TableS3_24hCA_reQTL_Models.xlsx)

Results from the 2df-model (tab 1), 1df-discrete (tab 2) and pseudobulk LME model (tab 3).

**Table S4. Results from reQTL analysis in PA.** (See Valencia_etal_TableS4_24hPA_reQTL_Models.xlsx)

Results from the 2df-model (tab 1), 1df-discrete (tab 2) and pseudobulk LME model (tab 3).

**Table S5. Results from reQTL analysis in MTB.** (See Valencia_etal_TableS5_24hMTB_reQTL_Models.xlsx)

Results from the 2df-model (tab 1), 1df-discrete (tab 2) and pseudobulk LME model (tab 3).

**Table S6. Results from reQTL analysis in IAV by cell type.** (See Valencia_etal_TableS6_IAV.xlsx)

Results from the 2df-model in B cells (tab 1), CD4+ T cells (tab 2), CD8+ T cells (tab 3), Monocytes (tab 4), and NK cells (tab 5).

**Table S7. Results from reQTL analysis in CA by cell type.** (See Valencia_etal_TableS7_24hCA.xlsx)

Results from the 2df-model in B cells (tab 1), CD4+ T cells (tab 2), CD8+ T cells (tab 3), NK cells (tab 4), Dendritic cells (DC) (tab 5), and Monocytes (tab 6).

**Table S8. Results from reQTL analysis in PA by cell type.** (See Valencia_etal_TableS8_24hPA.xlsx)

Results from the 2df-model in B cells (tab 1), CD4+ T cells (tab 2), CD8+ T cells (tab 3), NK cells (tab 4), Dendritic cells (DC) (tab 5), and Monocytes (tab 6).

**Table S9. Results from reQTL analysis in MTB by cell type.** (See Valencia_etal_TableS9_24hMTB.xlsx)

Results from the 2df-model in B cells (tab 1), CD4+ T cells (tab 2), CD8+ T cells (tab 3), NK cells (tab 4), Dendritic cells (DC) (tab 5), and Monocytes (tab 6).

**Table S10. Test of heterogeneous effect for the reQTL effect in IAV, CA, PA, and MTB** (See Valencia_etal_TableS10_IAV_CA_PA_MTB.xlsx)

Test of heterogeneity of each of the interaction terms (perturbation discrete=QE.Discrete and perturbation score=QE.Score) between cell types using the Cochran’s Q-test (QE). We consider heterogenous effect those with QE test Q value<0.10. IAV (tab 1), CA (tab 2), PA (tab 3), and MTB (tab 4).

**Table S11. Colocalization analysis on the non-perturbated and perturbated states in IAV, CA, MTB, and PA (See** Valencia_etal_TableS11_STIMs_Coloc_Discrete.xlsx)

Results from the colocalization analysis between the eQTL in the non-perturbated (NP) and perturbated (P) states and the GWAS locus. We used summary statistics of 9 GWAS on immune traits (Type 1 diabetes, Multiple sclerosis, Asthma, Rheumatoid arthritis, systemic lupus erythematosus, inflammatory bowel disease, Crohn’s disease, ulcerative colitis, psoriasis) and 3 GWAS on non-immune traits (coronary artery disease, type 2 diabetes, depression). We ran coloc with the default parameters and defined colocalization as those loci with a posterior probability H_4_>0.5. IAV (tab 1), CA (tab 2), PA (tab 3), and MTB (tab 4).

**Table S12. Colocalization analysis at different levels of the perturbation score in IAV, CA, MTB, and PA (See** Valencia_etal_TableS12_STIMs_Coloc_Tertiles.xlsx)

Results from the colocalization analysis between the eQTL at tertiles of the perturbation score within NP (NP_T1_-NP_T3_) and P (P_T1_-P_T3_) states and the GWAS locus. We used summary statistics of 9 GWAS on immune traits (Type 1 diabetes, Multiple sclerosis, Asthma, Rheumatoid arthritis, systemic lupus erythematosus, inflammatory bowel disease, Crohn’s disease, ulcerative colitis, psoriasis) and 3 GWAS on non-immune traits (coronary artery disease, type 2 diabetes, depression). We ran coloc with the default parameters and defined colocalization as those loci with a posterior probability H_4_>0.5. IAV (tab 1), CA (tab 2), PA (tab 3), and MTB (tab 4).

### Supplementary figures

**Figure S1.**
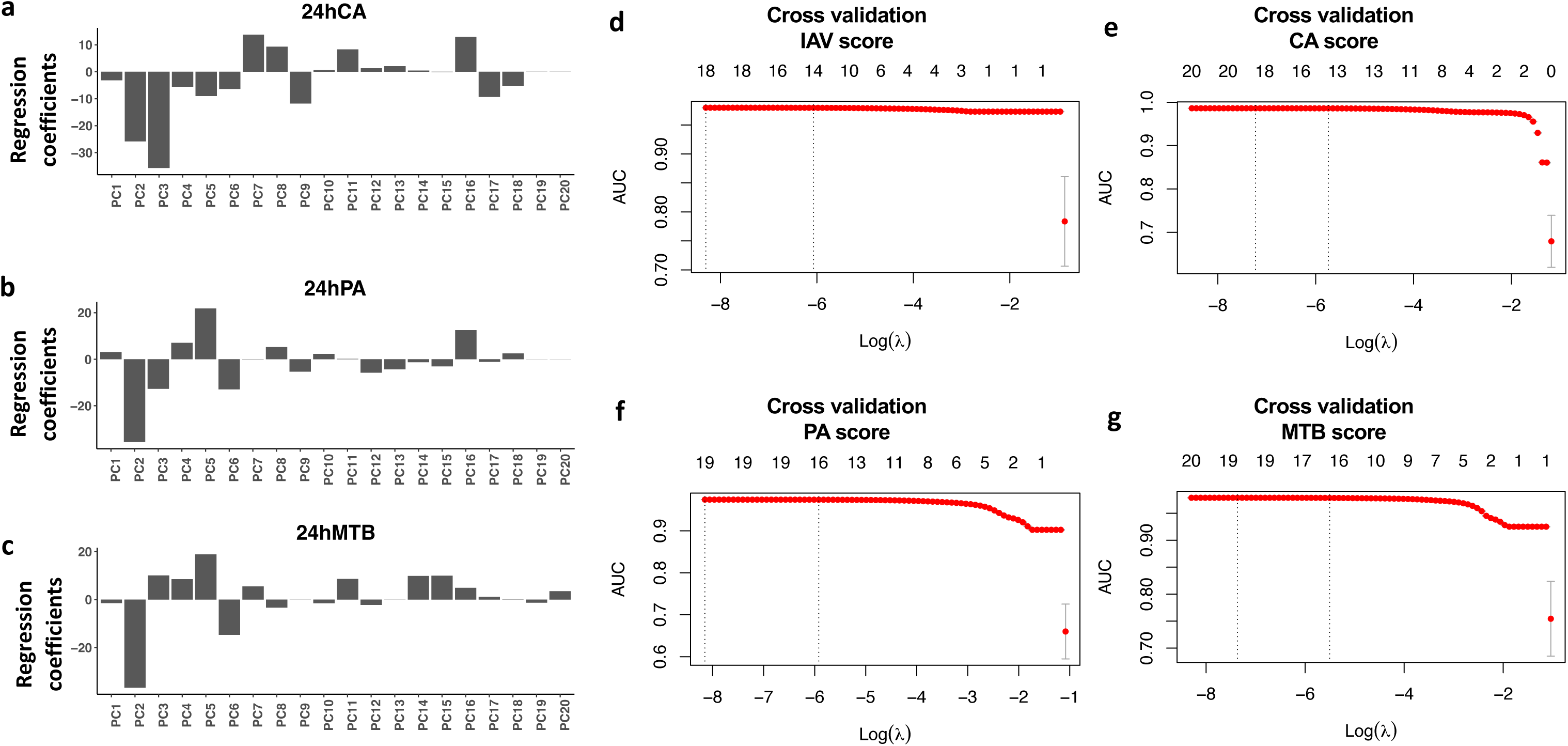
Modeling the perturbation score. We generated the continuous perturbation score in the CA, PA and MTB experiments. (a-c). The plot shows the regression coefficients of the penalized logistic regression in each of the perturbation experiments. (d-g) Plots showing the cross-validation results for each of the perturbation scores. Dotted lines shows the penalizations factor (𝜆) that resulted in the most regularized model and 𝜆 that gives the model with minimum mean cross validation error.

**Figure S2.**
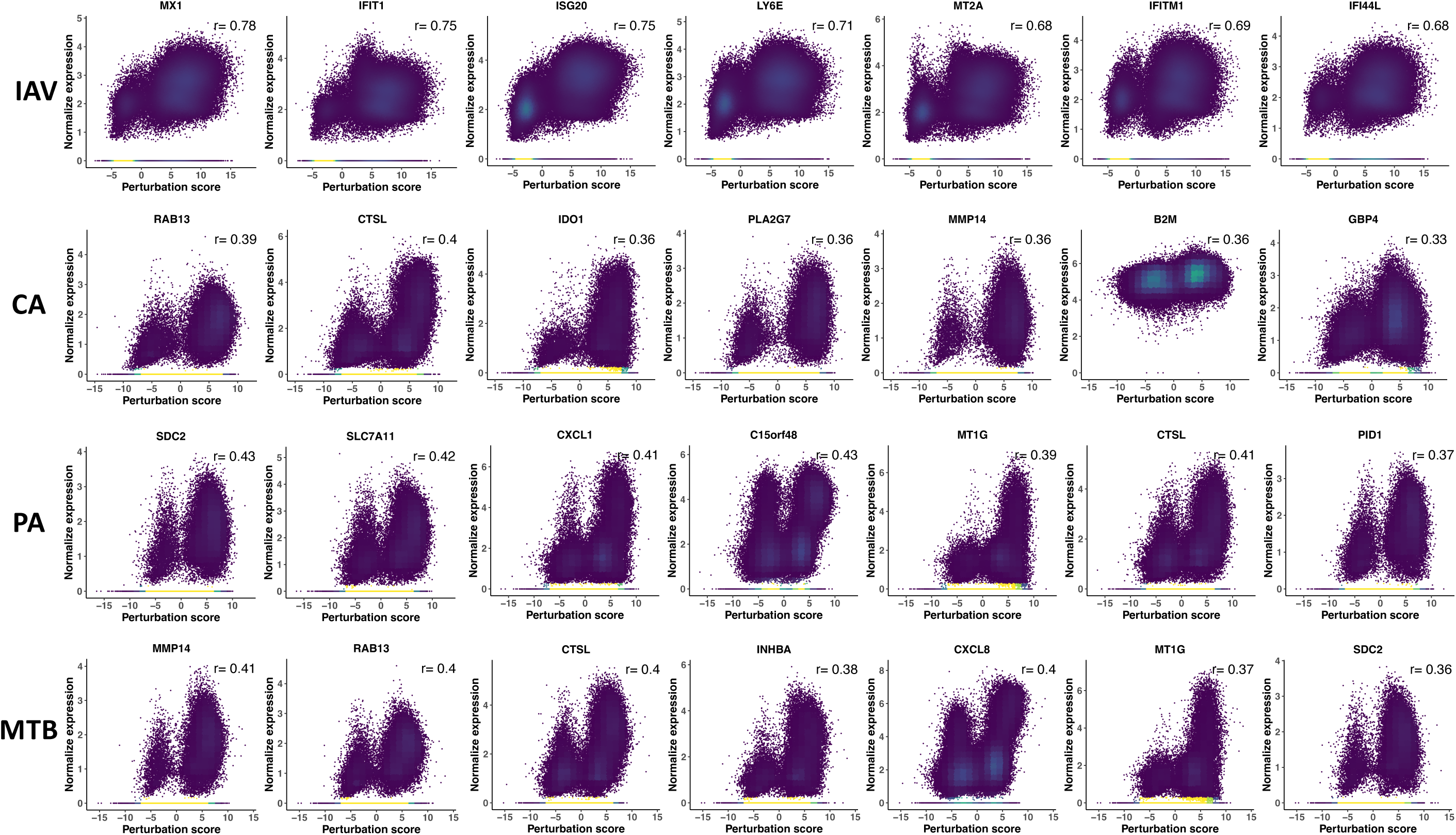
Top genes correlated with the perturbation score. Scatter plots showing the correlation of each perturbation score and the expression of top correlated genes colored by cell density.

**Figure S3.**
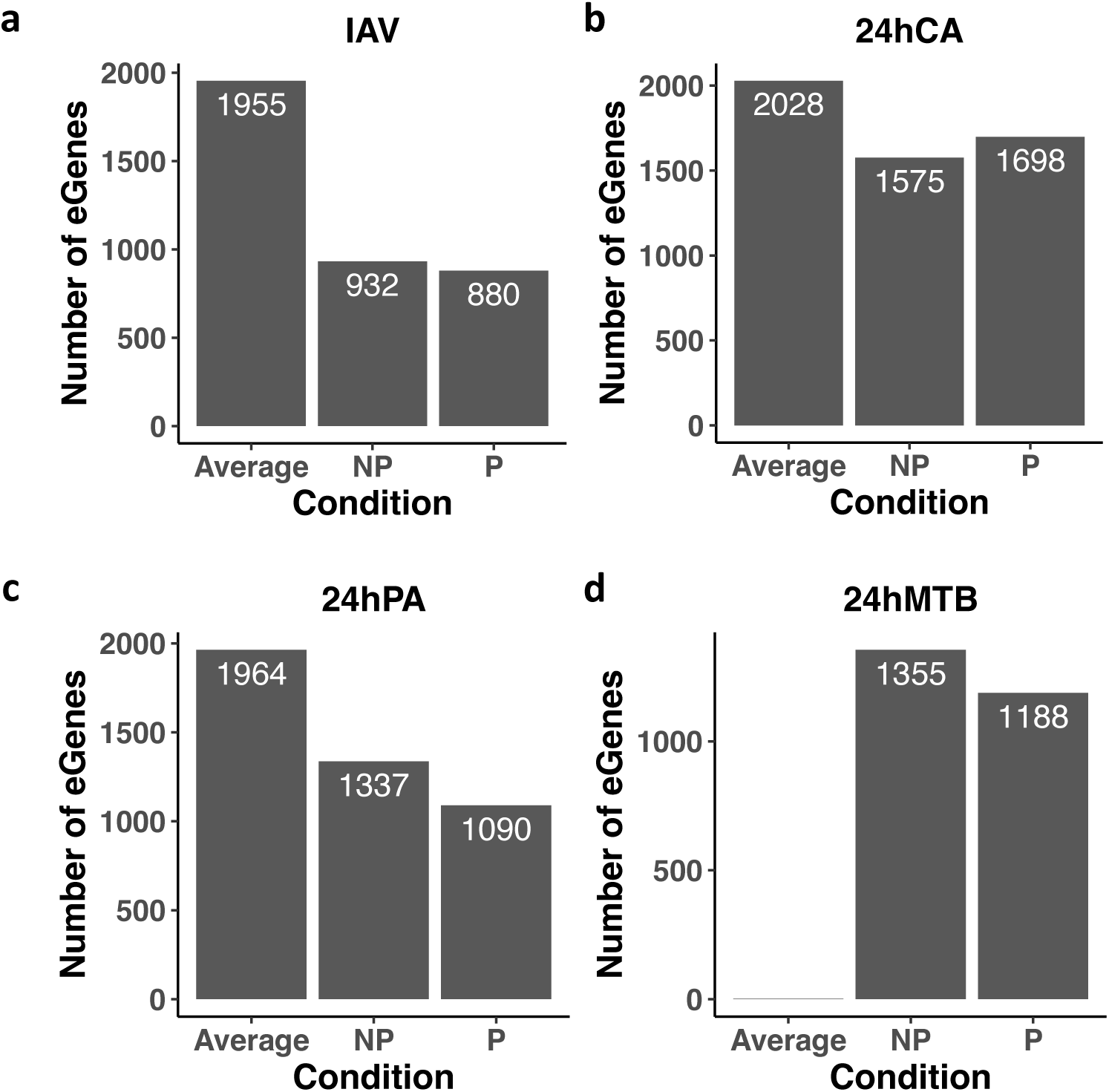
Set of eQTL Prioritized on each perturbation experiment. Barplots shows the number of significant (Q value <0.10) eQTL detected on bulk expression profiles for the non-perturbated (NP), perturbated (P) and the average of the NP and P conditions (**METHODS**).

**Figure S4.**
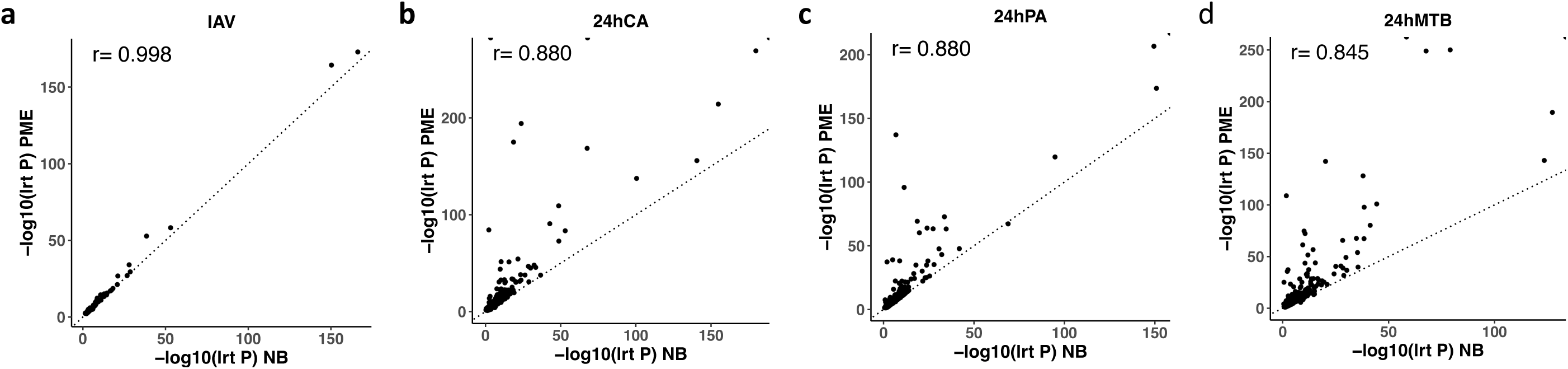
Comparison of the p values between PME and Negative binomial reQTL models. Scatter plot shows good agreement (Pearson correlation r) between the likelihood ratio test P (LRT P) from the single cell Poisson Mix Effect (PME) model (2df-model) and the LRT P from an equivalent negative binomial model (**METHODS**). Each dot represents a reQTL that was significant in the PME 2df-model.

**Figure S5.**
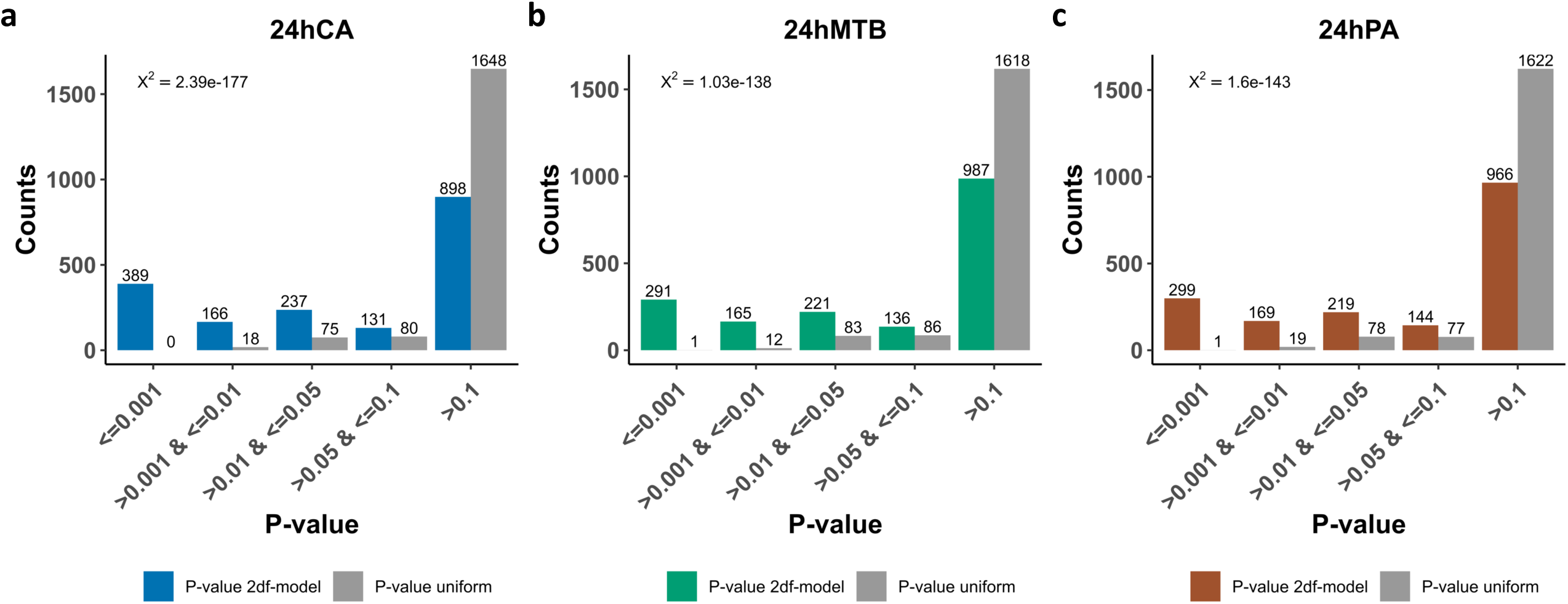
Enrichment of significant interaction among previously reported eQTLs. Results from our secondary analysis where we examined reQTL effects with the 2df-model on the set of 1,825 previously prioritized eQTLs from the Oelen et al 2022. (**METHODS**). X-axis indicates the number of reQTL with the specified LRT P value obstained on the 2df-model or under the uniform distribution (1,825 P values). X^2^ chi square test.

**Figure S6.**
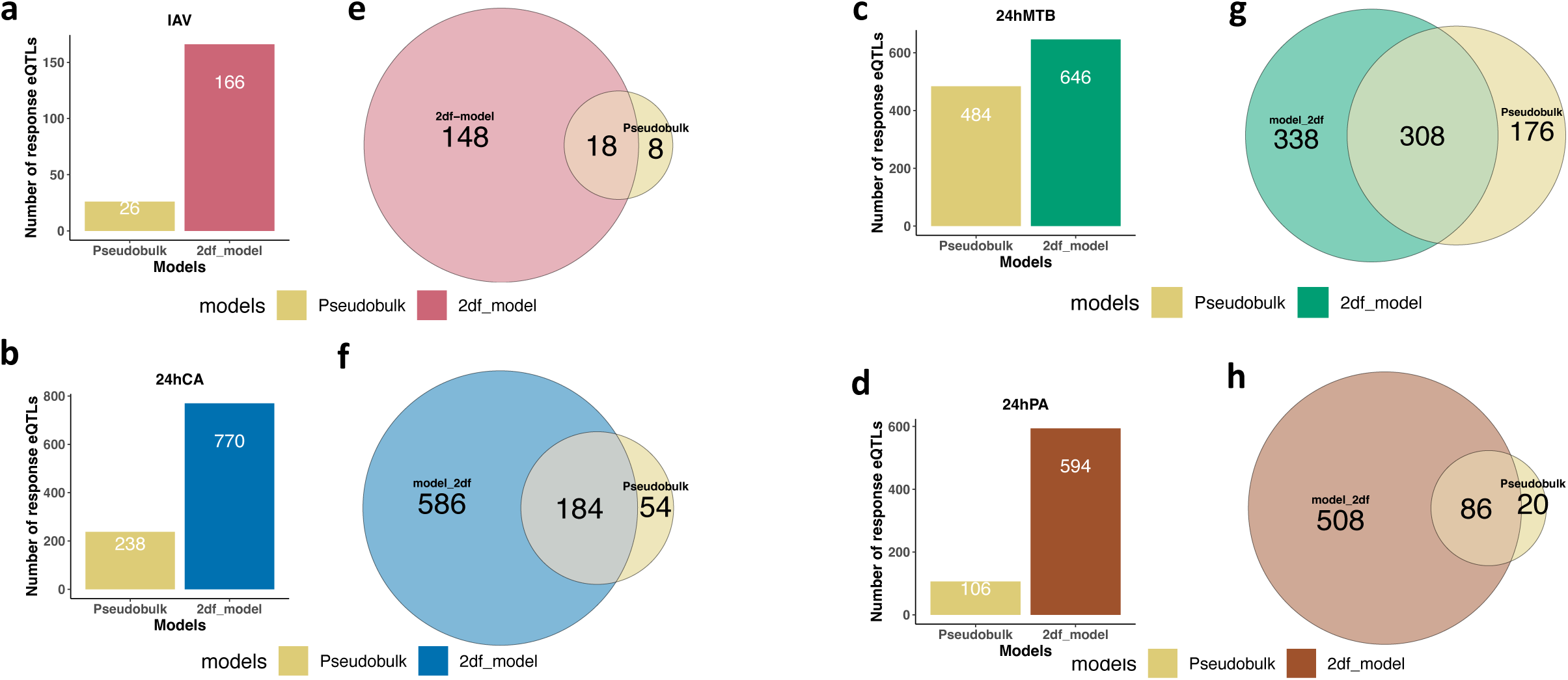
Mapping reQTL in single cells with the 2df-model have more power compared to a pseudobulk approach. a-d. Bar plot showing the number of reQTL detected by the single cell 2df-model and the LME-pseudobulk (**METHODS**) (e-h) Veen diagram showing the number of eGenes with reQTL specifically detected by each model and the number of overlapping eGenes with reQTL effects.

**Figure S7.**
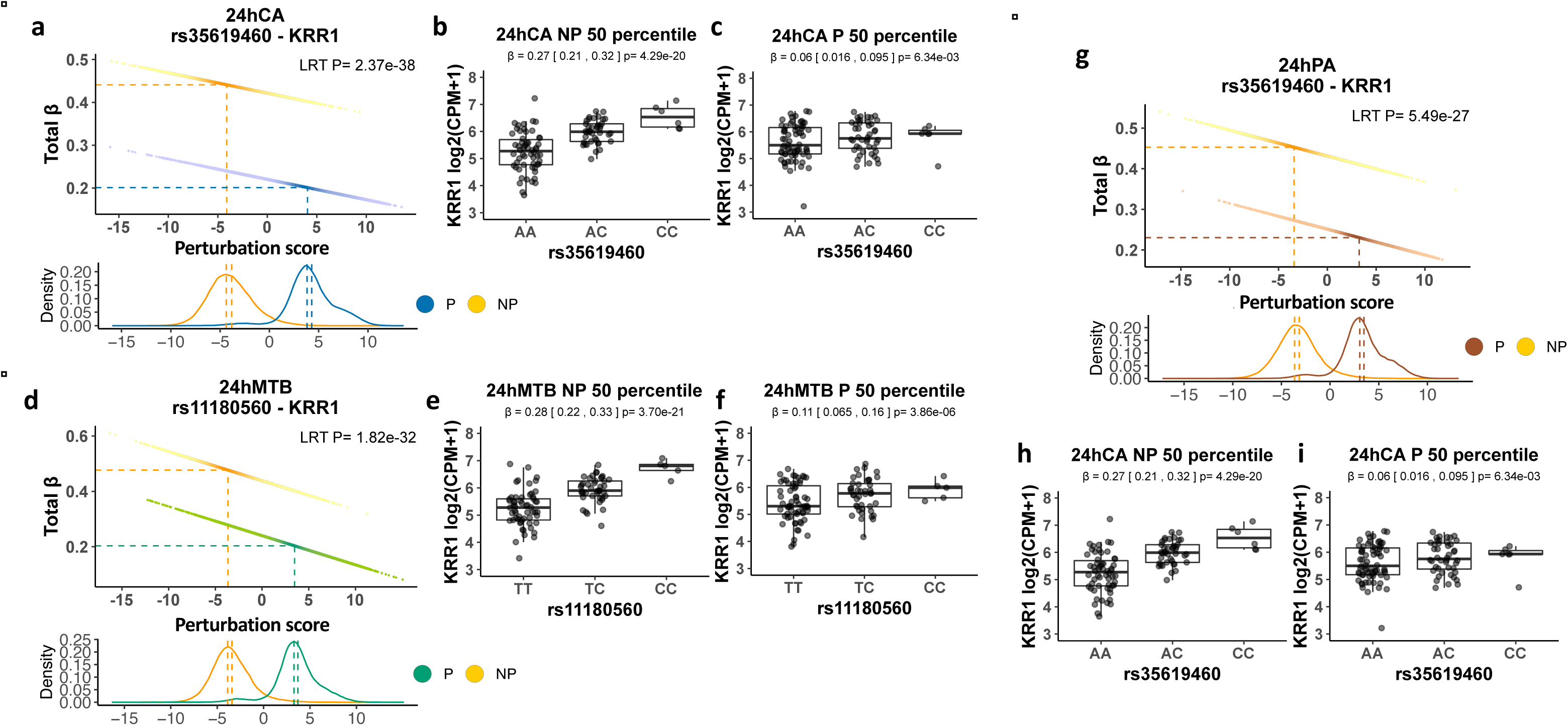
Decreased of the eQTL effect after perturbations in the KRR1 gene. Scatter plots showing the total eQTL effect for the gene KRR1 in the CA (a), MTB (d) and PA (g). Each dot represents the total beta in a cell. The color corresponds to the NP and P conditions. The dashed lines on the X axis indicates the perturbation score at the 50th percentile. The dashed line on the Y-axis indicates the total beta at the 50th percentile. The histogram below the scatter plot shows the density of the perturbation score color by NP and P conditions. Box plots showing the eQTL effect at the donor level for the eQTL effect for the gene KRR1 in the CA (b-c), MTB (e-f), and PA (h-i). Each boxplot represents the eQTL effect around the 50 percentiles on the NP and P conditions (**METHODS**).

**Figure S8.**
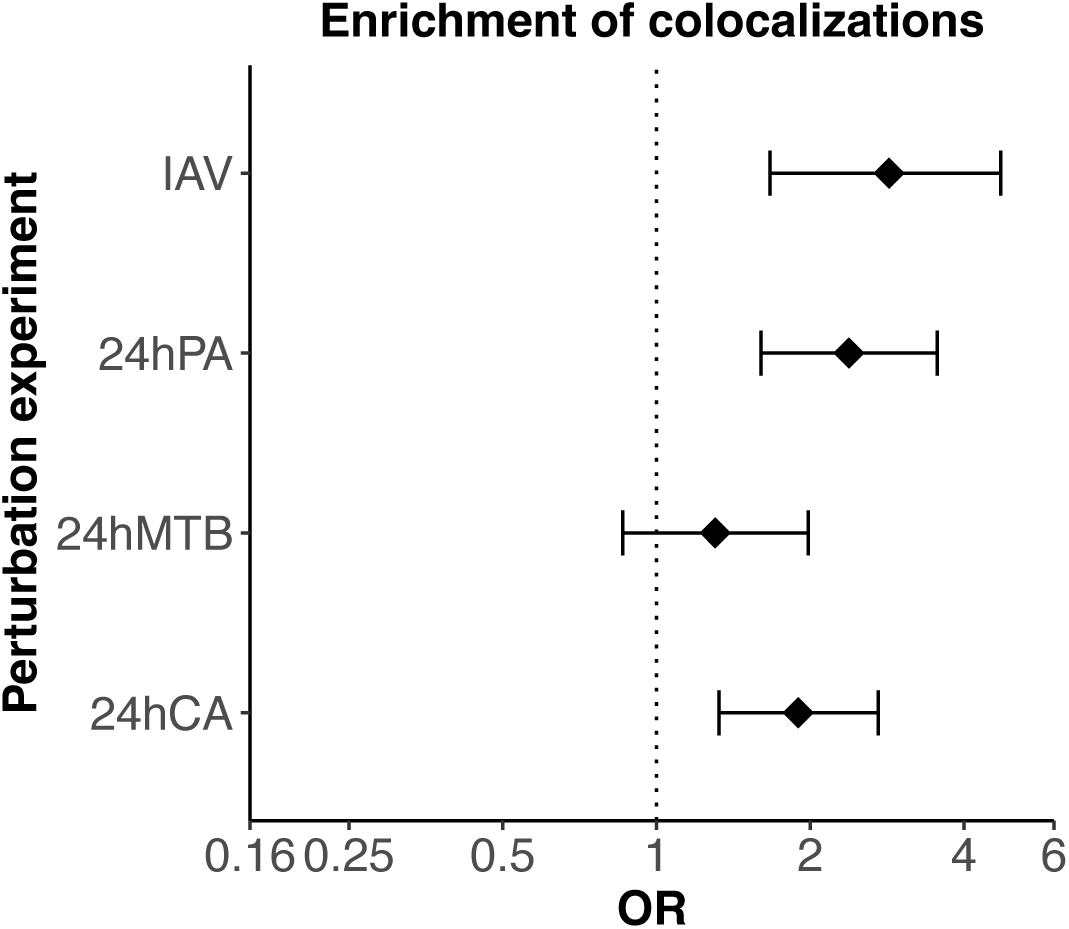
reQTL were enriched for colocalizations. Forrest plot shows that reQTL are more likely to colocalized compared to eQTL in the IAV, PA, and CA perturbation experiments (**METHOS**). OR=Odds ratio, error bars represented the 95% CI.

**Figure S9.**
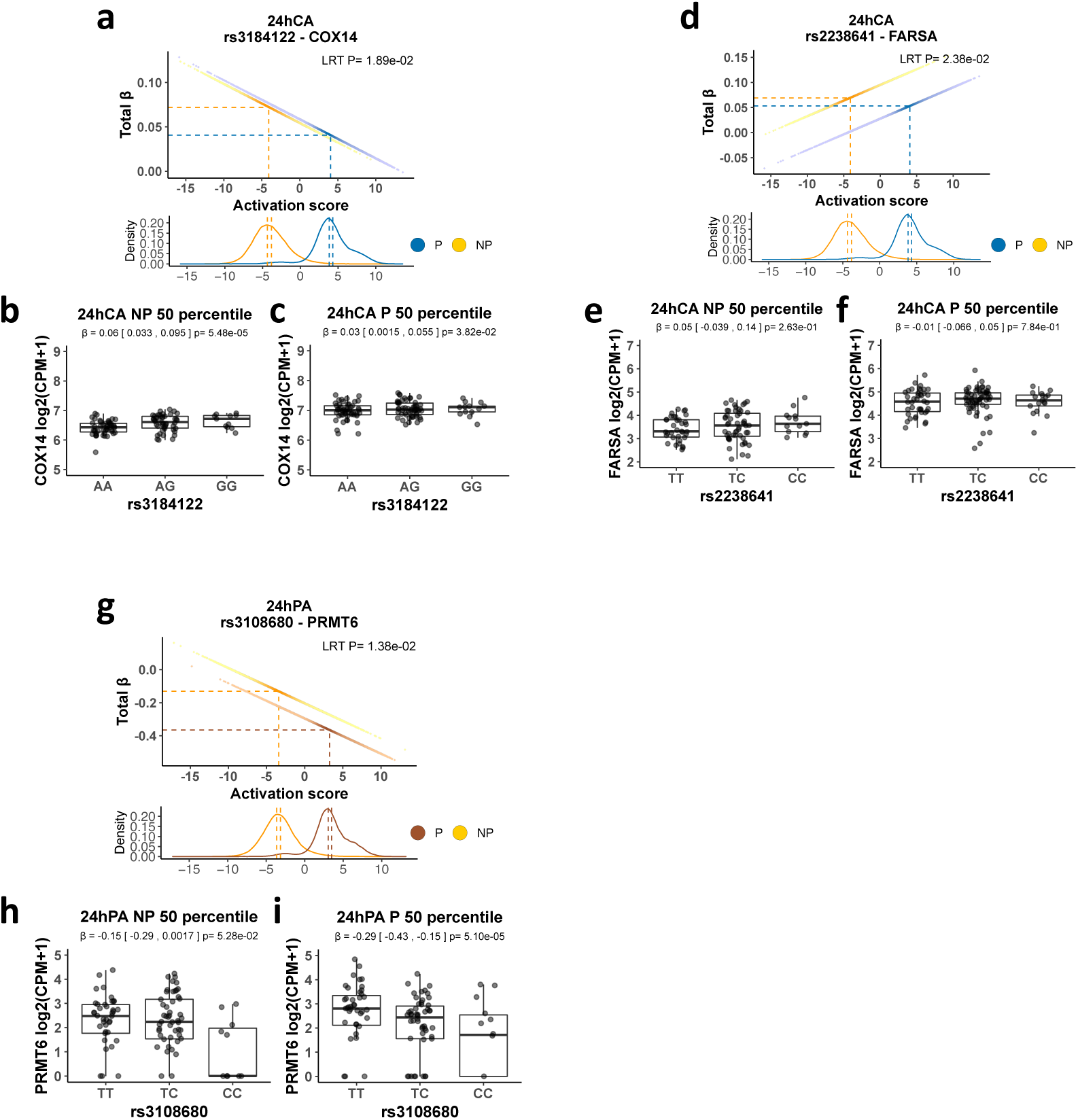
reQTL detected only by the 2df-model. a. Scatter plot showing the total eQTL effect of rs3184122 for the gene COX14 on each cell in the CA perturbation experiment. The color corresponds to the NP and P conditions. The dashed lines on the X axis indicates the perturbation score at the 50th percentile. The dashed line on the Y-axis indicates the total beta at the 50th percentile. b-c. Boxplot represents the eQTL effect of rs3184122 on COX14 in cells between the 45th to 55th percentiles within each, the NP and P state after aggregating the single cell gene count by donor. d. Total eQTL effect of rs2238641 for the gene FARSA on each cell in the CA perturbation experiment. e-f. Boxplot represents the eQTL effect of rs2238641 on FARSA in the NP and P states of the CA experiment. g. Total eQTL effect of rs3108680 for the gene PRMT6 on each cell in the PA perturbation experiment. h-i. Boxplot represents the eQTL effect of rs3108680 on PRMT6 the NP and P states of the PA experiment.

**Figure S10.**
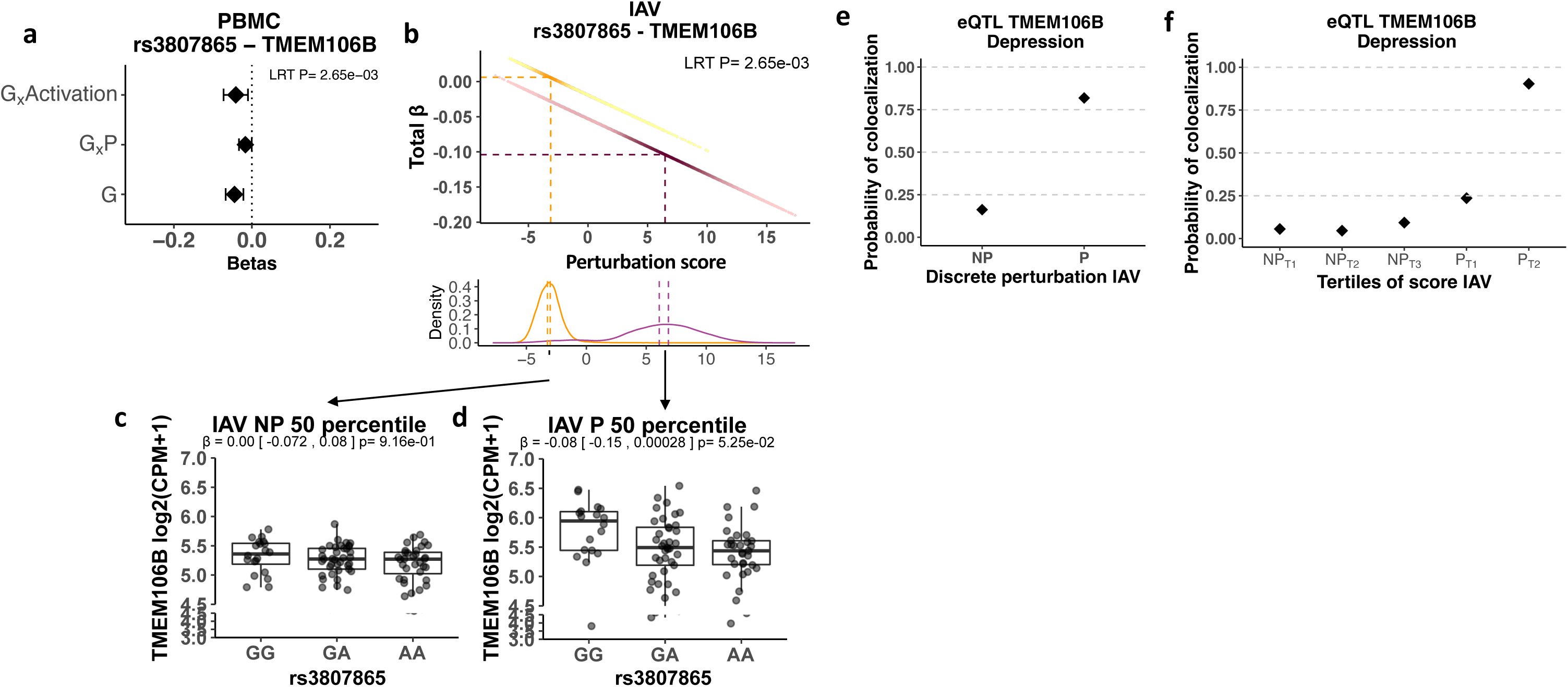
Increase of eQTL effect for TMEM106B in the IAV experiment. reQTL rs3807865 for the gene TMEM106B in the IAV perturbation experiment had perturbation-specific colocalization with depression. a. Forrest plot shows the main genotype (G) and interaction effects (G_x_Score, G_x_Discrete) from the 2df-model in PBMC. b. Scatter plot showing the total eQTL effect of rs3807865 for TMEM106B on each cell. The color corresponds to the NP and P conditions. The dashed lines on the X axis indicates the perturbation score at the 50th percentile. The dashed line on the Y-axis indicates the total beta at the 50th percentile. c-d. Boxplot represents the eQTL effect in cells between the 45th to 55th percentiles within each, the NP and P state after aggregating the single cell gene count by donor. e. Probability of colocalization (**METHODS**) of eQTL signal in TMEM106B with GWAS for depression at NP and P discrete state. f. Probability of colocalization of eQTL signal in TMEM106B with GWAS for depression at tertiles of the perturbation score (**METHODS**) in the NP (NP_T1_-NP_T3_) and P (P_T1_-P_T3_) cells. G= genotype term, G_x_Discrete=Interaction term between G and discrete perturbation state, G_x_Score= Interaction term between G and the continuous perturbation score.

**Figure S11.**
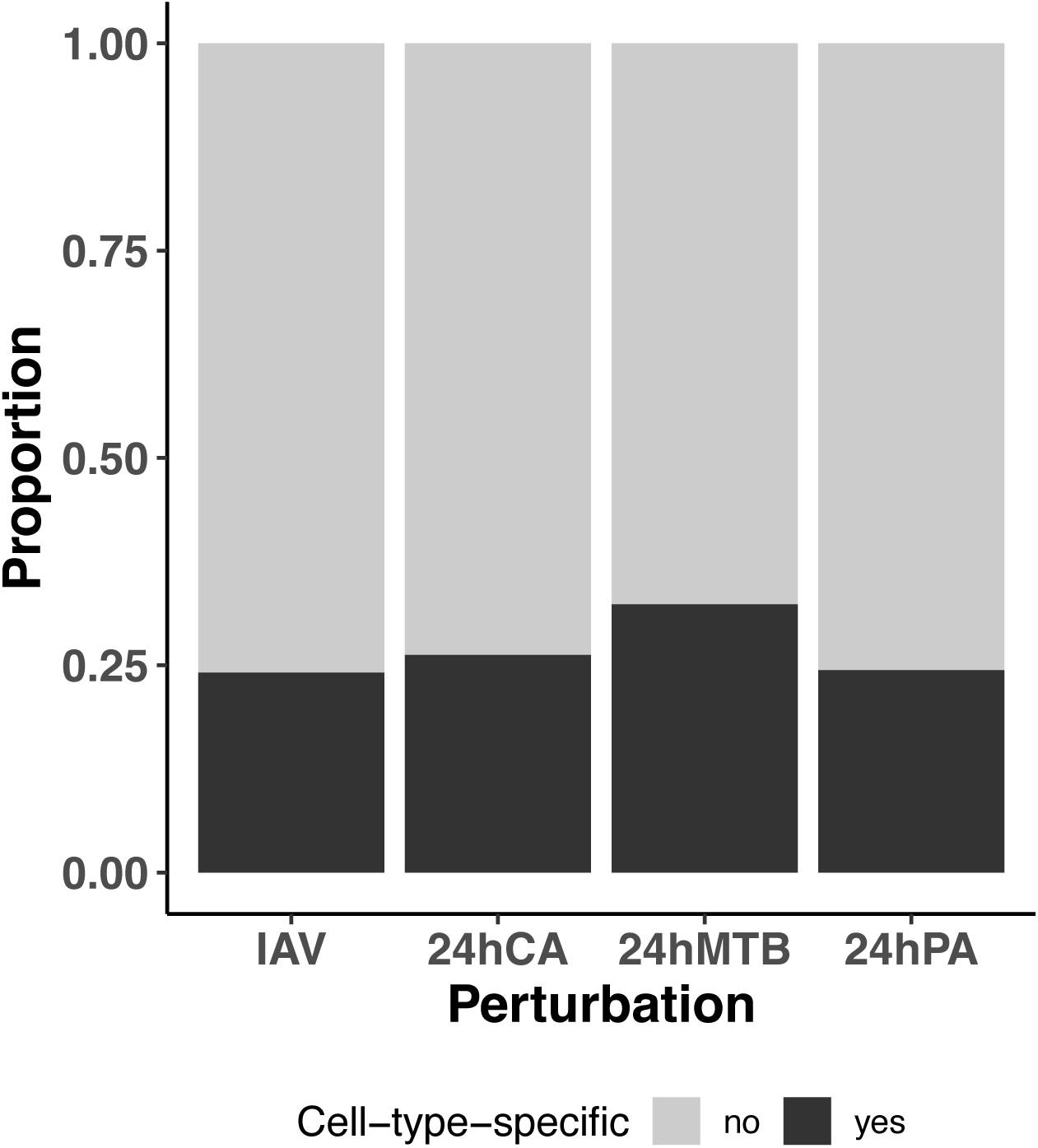
Proportion of reQTL with cell-type-specific effects. Barplot showing the proportion of reQTL with cell-type-specific effects on each of the perturbation experiments (**METHODS**).

**Figure S12.**
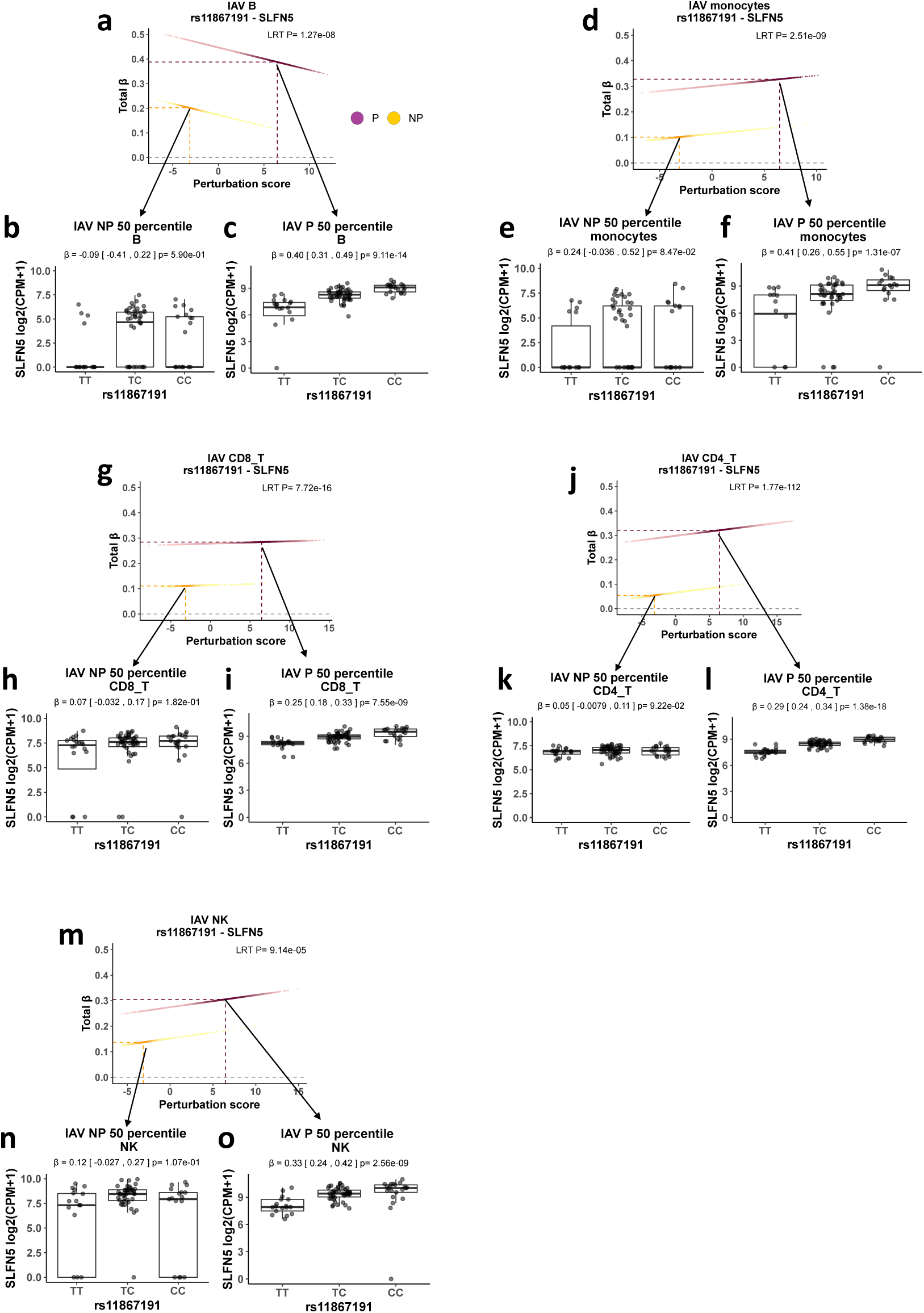
reQTL rs11867191 for SLFN5 have cell-type-specific effects in the IAV experiment. Scatter plot showing the total eQTL effect of rs11867191 for the gene SLFN5 on each cell in the IAV perturbation experiment. The color corresponds to the NP and P conditions. The dashed lines on the X axis indicates the perturbation score at the 50th percentile. The dashed line on the Y-axis indicates the total beta at the 50th percentile. Boxplot represents the eQTL effect of rs11867191 for the gene SLFN5 in cells between the 45th to 55th percentiles within each, the NP and P state after aggregating the single cell gene count by donor. a-c. B cells, d-f. Monocytes. g-i. CD8 T cells, j-l. CD4 T cells, m-o. Natural killer (NK) cells.

**Figure S13.**
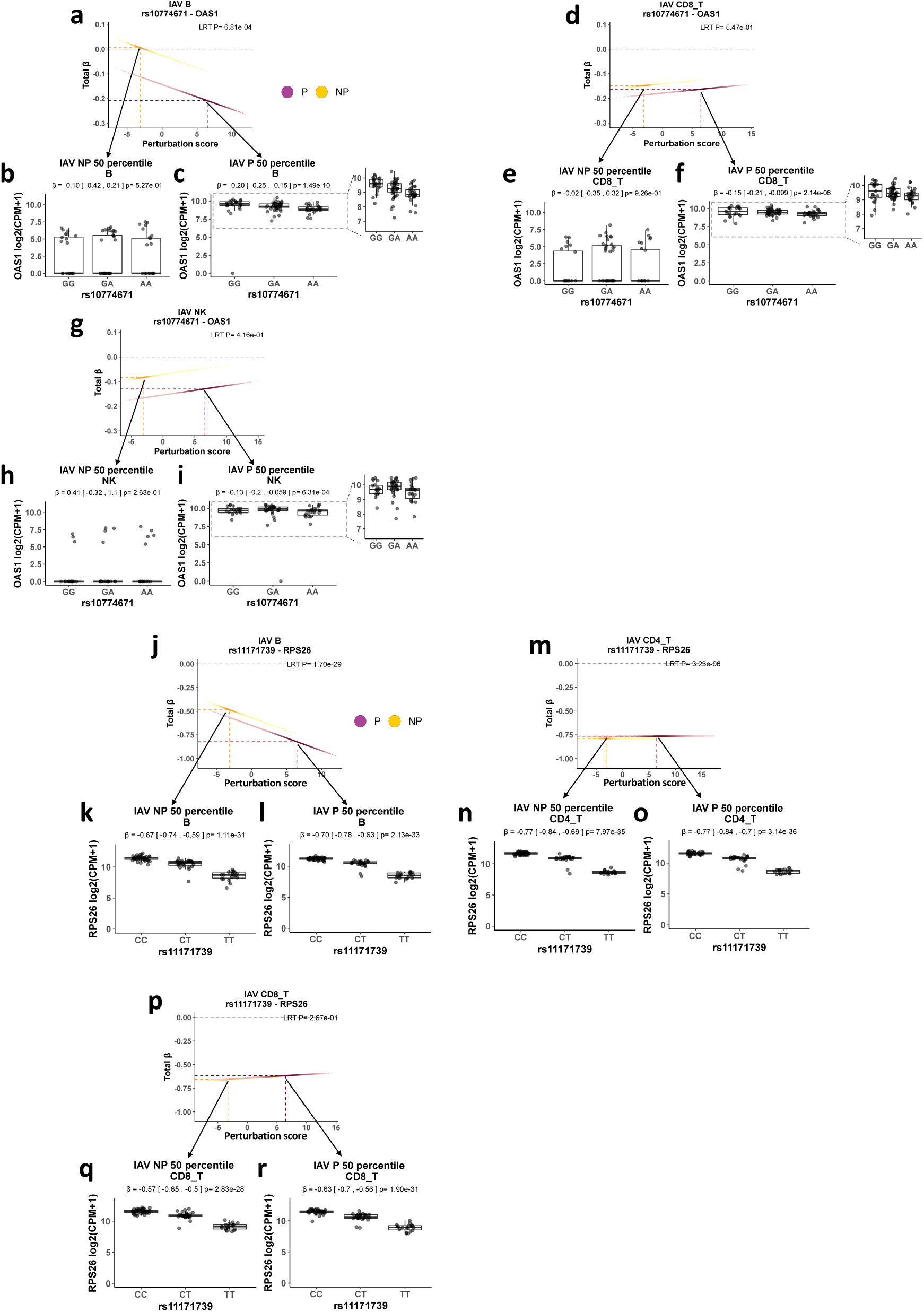
reQTL with cell-type-specific effects in the IAV experiment. Scatter plot showing the total eQTL effect on each cell in the IAV perturbation experiment. The color corresponds to the NP and P conditions. The dashed lines on the X axis indicates the perturbation score at the 50th percentile. The dashed line on the Y-axis indicates the total beta at the 50th percentile. Boxplot represents the eQTL effect in cells between the 45th to 55th percentiles within each, the NP and P state after aggregating the single cell gene count by donor. a-c. reQTL rs10774671 for the gene OAS1 in B cells. d-f. reQTL rs10774671 for the gene OAS1 in CD8 T cells. g-i. reQTL rs10774671 for the gene OAS1 in Natural Killer (NK) cells. Boxplots b, c ,e, f, h, and i. the area delimited by the dashed box was magnified. j-l. reQTL rs11171739 for the gene RPS26 in B cells. m-o. reQTL rs11171739 for the gene RPS26 in CD4 T cells. p-r. reQTL rs11171739 for the gene RPS26 in CD8 T cells.

**Figure S14.**
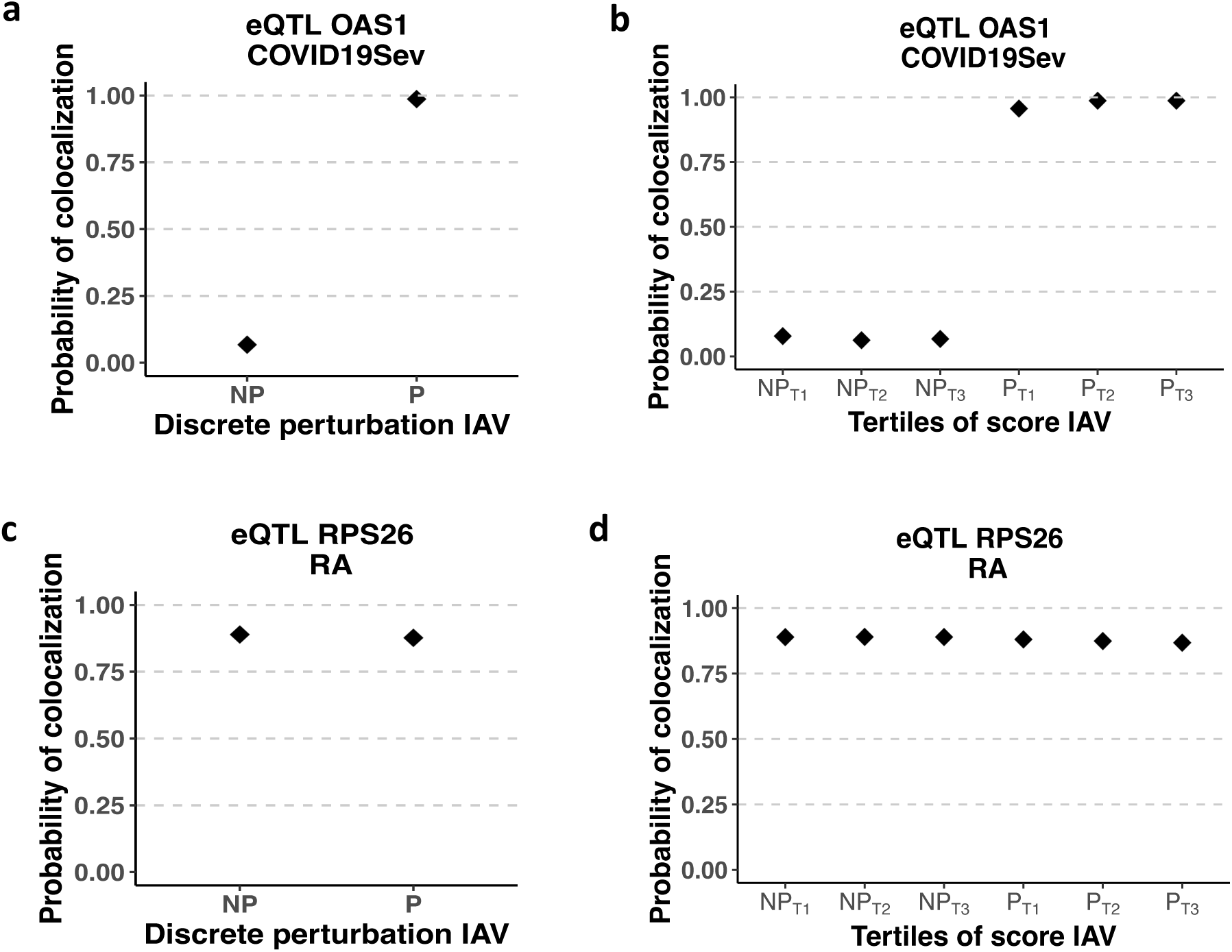
Colocalization of reQTL in the IAV experiment. Probability of colocalization (**METHODS**) of (a) eQTL signal in OAS1 with GWAS for severe Covid-19 at NP and P discrete state. b. Probability of colocalization of eQTL signal in OAS1 with GWAS for severe Covid-19 at tertiles of the perturbation score (**METHODS**) in the NP (NP_T1_-NP_T3_) and P (P_T1_-P_T3_) states. c Probability of colocalization of eQTL signal in RPS26 with GWAS for rheumatoid arthritis (RA) at NP and P discrete state. d. Probability of colocalization of eQTL signal in RPS26 with GWAS for RA at tertiles of the perturbation score in the NP (NP_T1_-NP_T3_) and P (P_T1_-P_T3_) states.

**Figure S15.**
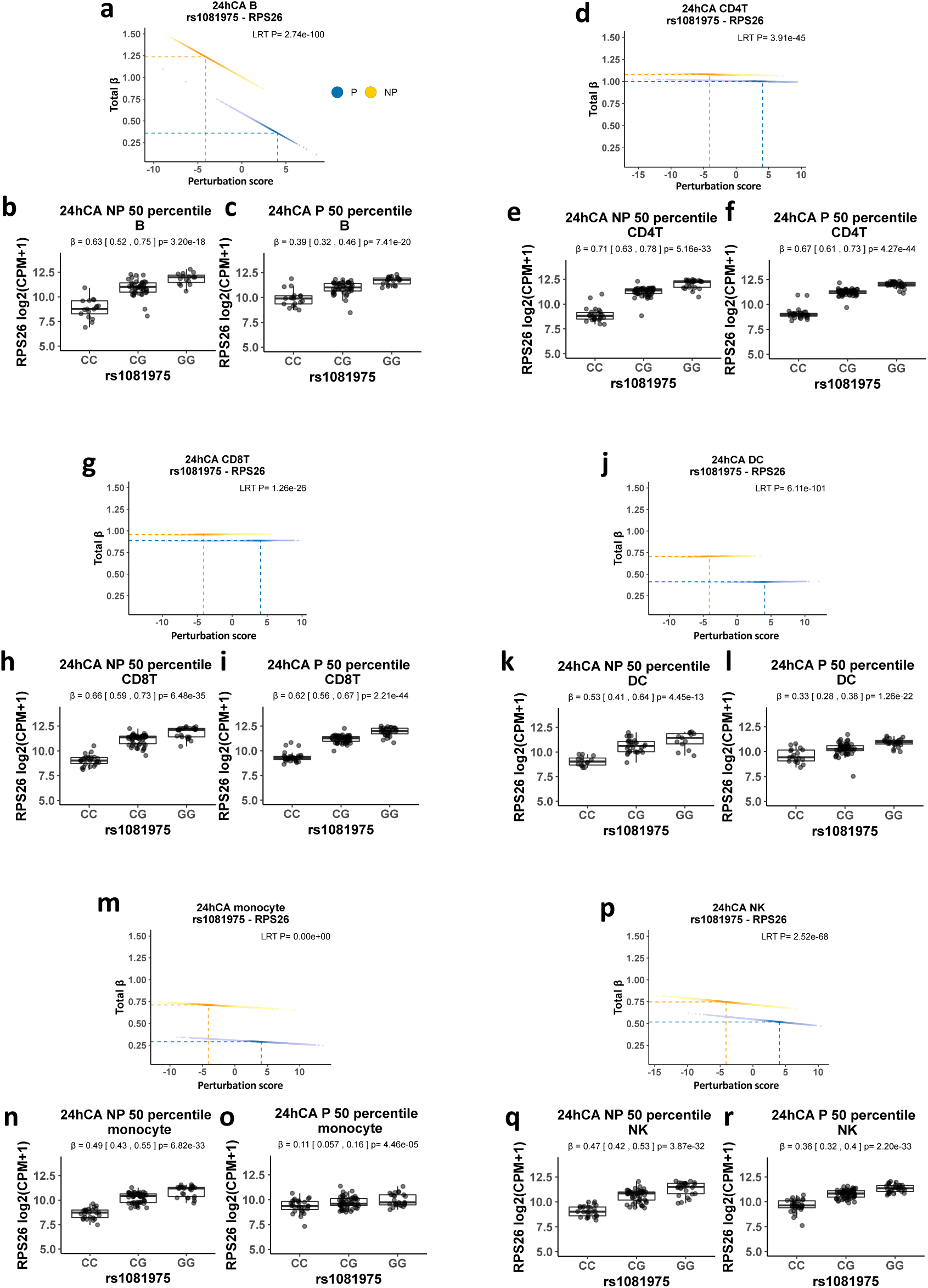
reQTL rs1081975 for RPS26 with cell-type-specific effects in the CA experiment. Scatter plot showing the total eQTL effect on each cell. The color corresponds to the NP and P conditions. The dashed lines on the X axis indicates the perturbation score at the 50th percentile. The dashed line on the Y-axis indicates the total beta at the 50th percentile. Boxplot represents the eQTL effect in cells between the 45th to 55th percentiles within each, the NP and P state after aggregating the single cell gene count by donor. a-c. reQTL rs1081975 for RPS26 in B cells. d-f. reQTL rs1081975 for RPS26 in CD4 T cells. g-i. reQTL rs1081975 for RPS26 in CD8 T cells. j-l. reQTL rs1081975 for RPS26 in Dendritic cells (DCs). m-o. reQTL rs1081975 for RPS26 in Monocytes. p-r. reQTL rs1081975 for RPS26 in Natural Killer (NK) cells..

**Figure S16.**
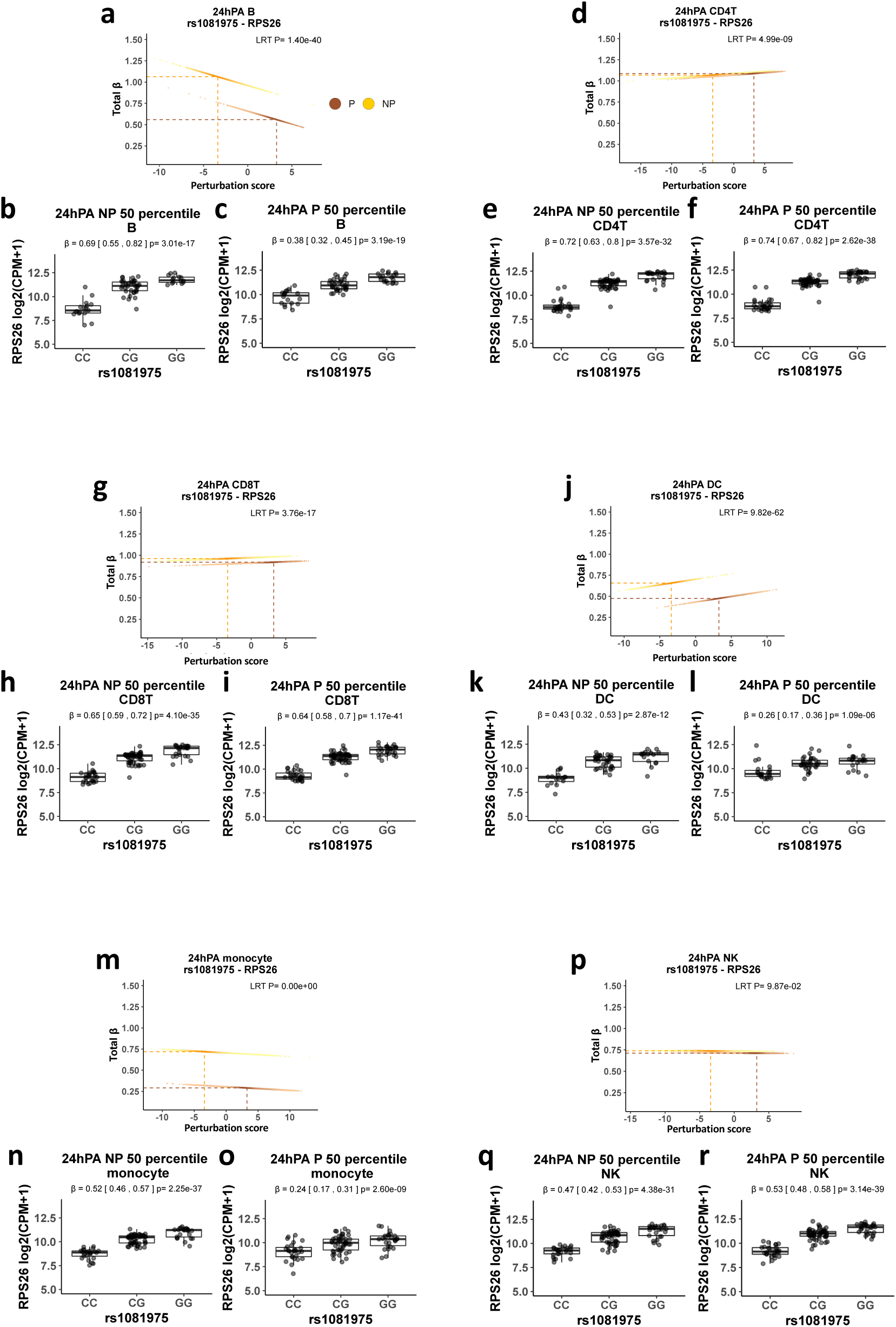
reQTL rs1081975 for RPS26 with cell-type-specific effects in the PA experiment. Scatter plot showing the total eQTL effect on each cell. The color corresponds to the NP and P conditions. The dashed lines on the X axis indicates the perturbation score at the 50th percentile. The dashed line on the Y-axis indicates the total beta at the 50th percentile. Boxplot represents the eQTL effect in cells between the 45th to 55th percentiles within each, the NP and P state after aggregating the single cell gene count by donor. a-c. reQTL rs1081975 for RPS26 in B cells. d-f. reQTL rs1081975 for RPS26 in CD4 T cells. g-i. reQTL rs1081975 for RPS26 in CD8 T cells. j-l. reQTL rs1081975 for RPS26 in Dendritic cells (DCs). m-o. reQTL rs1081975 for RPS26 in Monocytes. p-r. reQTL rs1081975 for RPS26 in Natural Killer (NK) cells..

**Figure S17.**
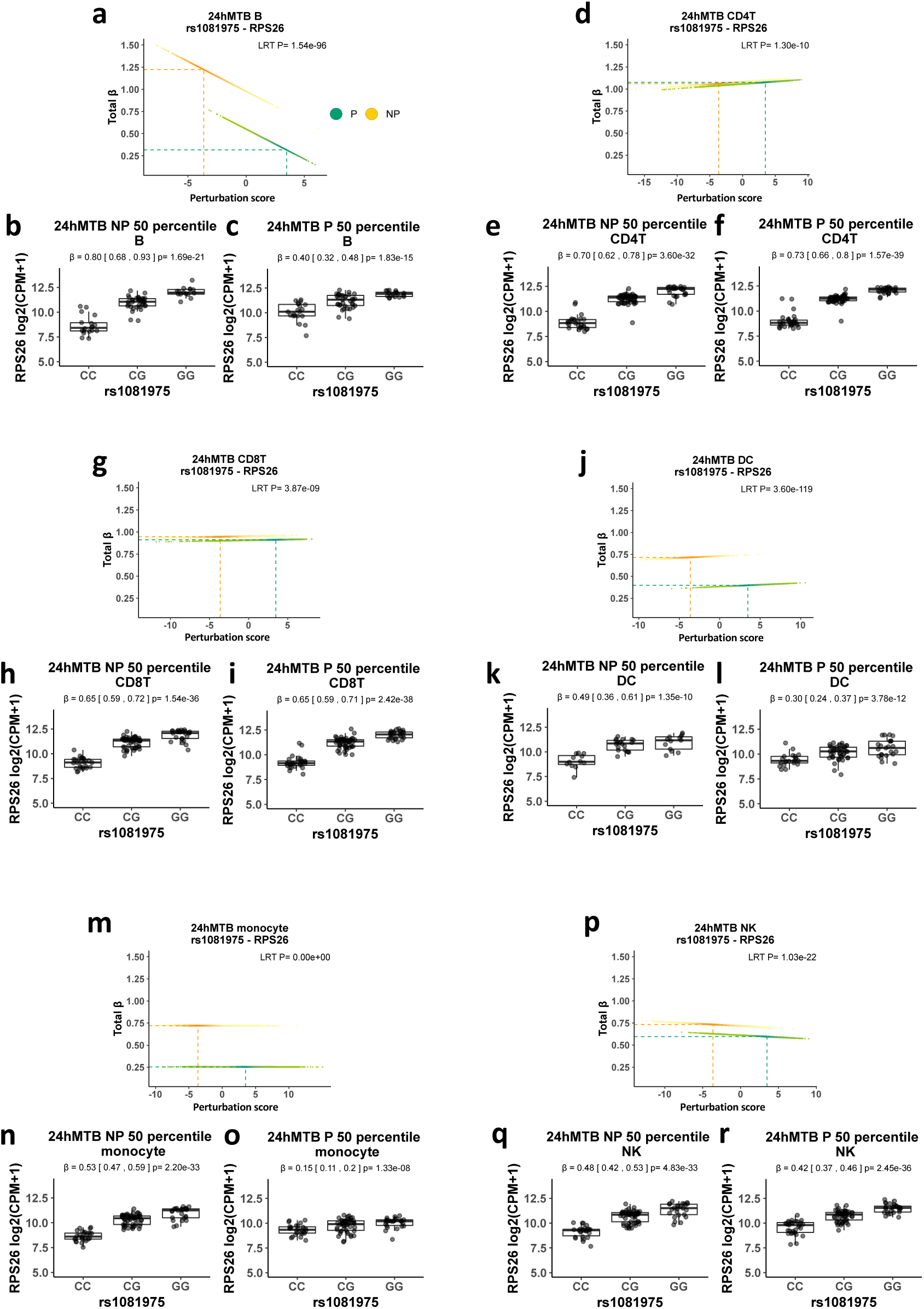
reQTL rs1081975 for RPS26 with cell-type-specific effects in the MTB experiment. Scatter plot showing the total eQTL effect on each cell. The color corresponds to the NP and P conditions. The dashed lines on the X axis indicates the perturbation score at the 50th percentile. The dashed line on the Y-axis indicates the total beta at the 50th percentile. Boxplot represents the eQTL effect in cells between the 45th to 55th percentiles within each, the NP and P state after aggregating the single cell gene count by donor. a-c. reQTL rs1081975 for RPS26 in B cells. d-f. reQTL rs1081975 for RPS26 in CD4 T cells. g-i. reQTL rs1081975 for RPS26 in CD8 T cells. j-l. reQTL rs1081975 for RPS26 in Dendritic cells (DCs). m-o. reQTL rs1081975 for RPS26 in Monocytes. p-r. reQTL rs1081975 for RPS26 in Natural Killer (NK) cells..

